# PEG400 regulates Falcipain 2 activity through an unprecedented allosteric mechanism

**DOI:** 10.1101/2025.10.29.685357

**Authors:** Bikram Nath, Subhoja Chakraborty, Sampa Biswas

## Abstract

The malarial parasite *Plasmodium falciparum* cleaves host hemoglobin by cascade of proteolytic enzymes. The cysteine protease, Falcipain-2 (FP2) plays an essential role in the process and important for parasite survival, making it a potential drug target. However, similarities with host cysteine cathepsins hamper selective inhibition, thus necessitates detailed structural and functional characterizations of FP2. The present study uncovers a novel regulatory role of polyethylene glycol 400 (PEG400) on FP2 activity. PEG400 inhibits FP2 activity on small peptide substrate and azo-casein, while enhancing hemoglobin degradation, rendering a dual effect on FP2 catalysis. A mixed-type of inhibition has been observed for PEG400 against small peptide substrate of FP2, consistent with binding of PEG400 to catalytic cleft, confirmed by fluorescence quenching and docking studies. Unlike typical nonspecific PEG-protein interactions, PEG400 adopts a fit within catalytic region of FP2 and partially overlaps with leupeptin binding sites, albeit with lower affinity. Computational analysis further identifies a novel allosteric binding pocket of PEG400, supported by in-silico mutagenesis and molecular dynamics simulation. This pocket exhibits minimal conservation in human cathepsins, suggesting selective potential. In contrast to this inhibitory role, biochemical assay reveals that PEG400 promotes haemoglobin proteolysis. Spectroscopic analyse suggests PEG400 alter hemoglobin structural dynamics to favour proteolysis. ENM based normal mode analysis reveals upon haemoglobin binding, PEG400 restricts FP2 hinge-bending motion, improves FP2-hemoglobin proximity, and simultaneously PEG400 is dislodged from the active site, thereby promoting proteolysis. The combined experimental and computational findings reveal a novel mechanism of FP2 regulation, opening new therapeutic avenues.

## INTRODUCTION

Malaria remains a significant global health concern, with 249 million cases and over 600,000 deaths having been reported globally in 2023 [1]. The severe and often fatal form of the disease is primarily caused by the *Plasmodium falciparum* parasite. The multifactorial dependence of malarial treatment on the parasite species, severity of the illness, the patient’s age and overall health along with the emergence of drug resistance to frontline Artemisinin-based combination therapies (ACTs) underscores the urgent need to identify novel drug targets and accelerate malarial chemotherapy with unique mechanisms of action [2].

Symptomatic exacerbation of *falciparum* malaria is executed by cysteine proteases called falcipains, which cleave host hemoglobin to facilitate parasite growth [3,4] among which Falcipain-2 (FP2) is pivotal in the early trophozoite stage. Structural analyses have revealed that the mature catalytic domain of FP2 is a single polypeptide chain of 241 amino acids that adopts a typical two-domain papain-like fold [Fig. 1a]. FP2 activity relies on specific cysteine and histidine residues forming a catalytic dyad [5]. The unique features of FP2 lie within a distinctive N-terminal projection crucial for protease folding and a β-hairpin loop involved in hemoglobin interaction [6]. Although a detailed mechanism of Hb-binding to FP2 is yet to be deduced, several crucial residues have been located on FP2, found to be indispensably involved in its interactions with its natural substrate Hb. FP2 reportedly prefers cleavage sites with Leu at the P2 position and Hb hydrolysis by malaria parasites proceeds with rapid cleavage by falcipains at multiple sites of Hb [7]. Excerpts from modelling studies suggest that the β-hairpin loop of FP2 mediates H-bond and/or Van der Waals interactions with Hb, via its residues Glu185 and Val187. Val187 may possibly have a role in stabilization of FP2-Hb complex via Van der Waals interactions [8]. Overall, a 4:1 ratio of enzyme to Hb has been validated in the FP2-Hb, where Hb being a tetramer, each subunit seemingly interacts with one molecule of the enzyme [9]. Notably, FP2 has been identified as a promising target for antimalarial therapy due to its seminal involvement in host hemoglobin degradation and in the rupture of host cell membrane facilitating parasite egress [10,11].

**Figure 1.**
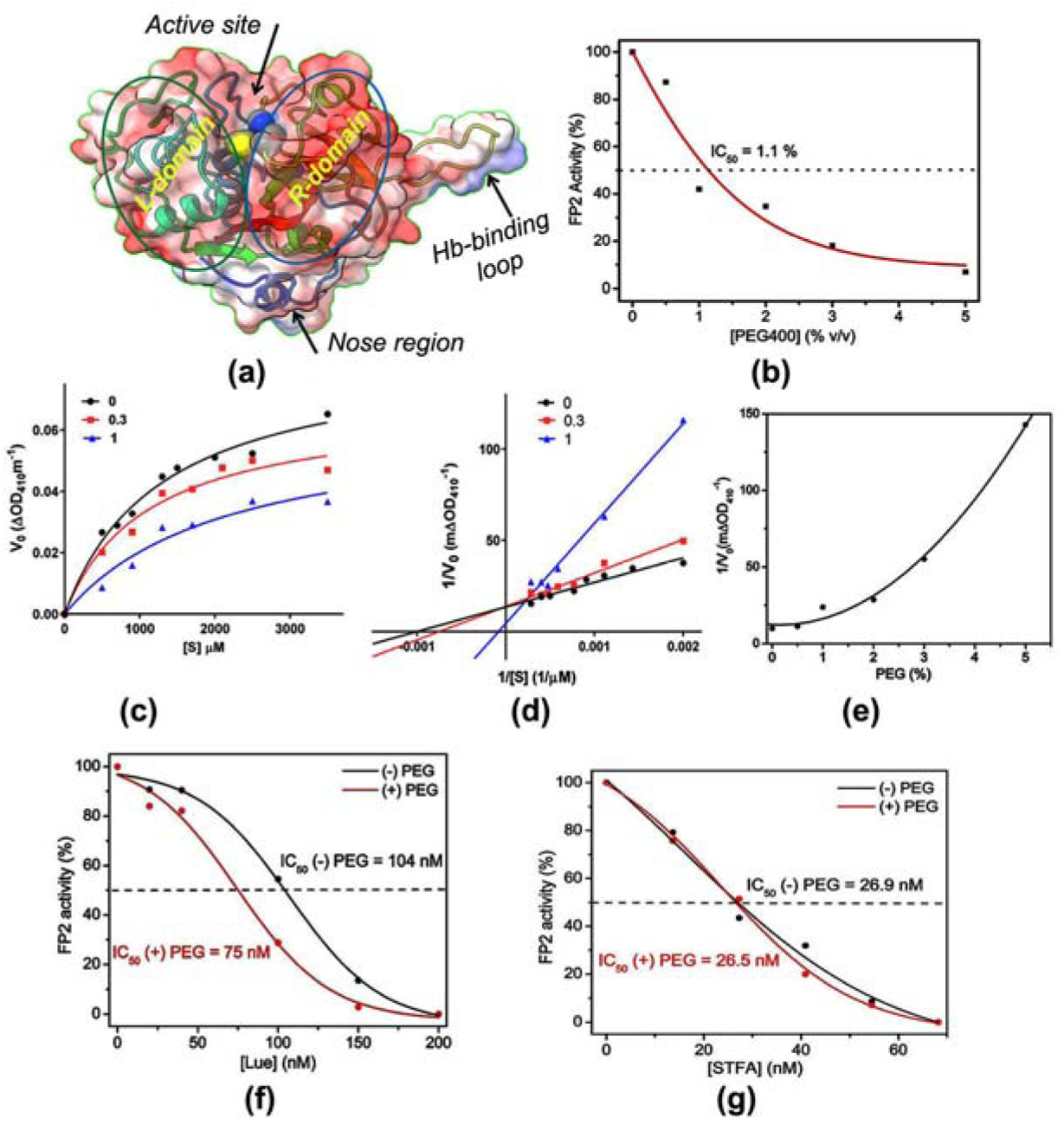
Structure and kinetics of FP2. (a) General structural organization of mature and active FP2 (b) IC_50_ calculation of PEG400 on FP2 using the substrate D-VLK-pNA (c) Initial velocity (V_0_) at different concentrations of D-VLK-pNA in presence of 0 - 1 % PEG 400 (d) Lineweaver-Burk (LB) plot using 0 – 1 % PEG400 and (e) Dixon plot using a fixed substrate concentration of 700 μM. IC_50_ of (f) leupeptin (Lue) and (g) Human stefin A (STFA) mediated inhibition of FP2 in absence and presence of 0.5 % PEG400 [denoted as (-) PEG and (+) PEG respectively].

Our recent research [6] has delved into the crystal structure of FP2 from the 3D7 strain of *Plasmodium falciparum*, shedding light on altered surface electrostatics and proposing a conformational active-inactive switch in its hemoglobinase activity. This structural insight has paved the way for potential drug design, highlighting new areas for intervention, such as allosteric binding sites similar to the orthologous human Cathepsin-K. Additionally, our finding [12] involves the efficient inactivation of FP2 by a human cystatin, stefin-A (STFA), offering promising strategy of developing potent inhibitors by targeting non-conserved secondary binding sites. However, inhibition of FP2 has been challenging because of the highly conserved active site with homologous host proteases-like cathepsins [13]. Therefore, understanding the regulatory mechanisms of FP2, in commensuration with its allosteric sites, has become important in this context.

The present study was initiated with understanding the effect of human Cathepsin-K-specific allosteric modifier chondroitin sulfate and heparin [14] on FP2 and found minimal effect on FP2 catalytic activity. The stable interaction of polyethylene glycol (PEG) with FP2 at ortho as well as allosteric sites, as observed in FP2 crystal structures from our group [6,11], encouraged us to investigate the role of PEG on the activity of FP2. Surprisingly, we observed polyethylene glycol, particularly PEG400, has a remarkable effect on the general proteolytic activity as well as specific hemoglobinase activity of the enzyme.

Within the cellular context, proteins navigate a complex environment comprising of diverse polymers and macromolecules of different sizes, shapes and concentrations that influence their behavior and regulate their activity [15–17]. *In vitro* studies of proteins in the presence of natural (DNA, proteins, carbohydrates, etc.) and synthetic (Ficoll, dextran, and PEGs) crowders have provided valuable insights into their interactions, and perpetual stability [18–20]. Polyethylene glycols are flexible and non-toxic polymers of varying sizes that allow their appropriate usage in different medical applications and as crowding agents to mimic intracellular environments in biological research [18].

Here, we have extensively explored the impact of polyethylene glycol with varying molecular weights (400, 4000 and 6000 Da) on FP2 activity against both small-molecule substrates and hemoglobin (Hb). Interestingly, PEG400 inhibited FP2 activity towards small substrates but enhanced Hb-degradation. In contrast, higher molecular weight PEGs displayed distinct behaviors as PEG 4000 showed minimal effect on FP2 activity, while PEG 6000 exhibited an impact similar to PEG400, albeit only at substantially higher concentrations. To ascertain whether PEG’s effect on Hb degradation was Hb-specific or extended to other macromolecules, we conducted azocasein assays. Intriguingly, azocasein behaved more akin to small molecular substrates than Hb, suggesting a unique interaction between FP2, PEG400 and Hb. Apart from the enzymatic activity; we also examined structural effects of PEGs on FP2, Hb and their complex through intrinsic Tryptophan fluorescence, far-UV CD and absorption spectroscopy. It is important to note that we have used a concentration range of PEG far below the threshold required for crowding effect, specifically for a low molecular weight polyethylene glycol as PEG400. Thus, this study mainly explores the effect of PEG as a potential ligand rather than a crowding agent.

To explore molecular insights into the structural rearrangement of FP2 underlying its altered activity in the presence of PEG, we decided to build reasonable models of each interacting partner of FP2, with the protease, through molecular modeling, docking studies, simulation and normal mode analysis. The combined results of targeted docking, simulation, cavity and ligand clusters analyses in corroboration with spectroscopic studies facilitate the identification of specific binding sites of PEG at both orthosteric and allosteric regions of FP2 with varying binding affinities. Notably, previous studies have reported that targeting the allosteric site of FP2 using synthetic small-molecule inhibitors and heme alters its structural organization with reduced activity [21–23]. However, the probable allosteric site of FP2, which we have identified in this study, is different from those reported earlier.

These findings provide novel insights into the probable *in-vivo* mechanism and regulatory aspects of FP2 in degradation of Hb and other substrates and highlight the effects of macromolecular ligands/ crowders in protein-protein interactions. The findings also gauge the occupancy of its multiple ligand-binding sites to aid in the development of targeted FP2 inhibitors with high specificity. Moreover, the observed interactions of PEG with multiple FP2 regions can provide a plausible framework for development of polymer-based drug delivery to targeted domains.

## RESULTS

### FP2 Inhibition

#### PEG400 as potential inhibitor

##### Identification

In lieu of the identification of allostery in papain-like proteases such as human cathepsins B, K and FP-2 itself, with outlined regulatory roles of heparin and other glycosaminoglycans [24–29] as well as our earlier work [30] and FP2 crystal structures analyses [6,12] identifying PEG molecules bound to the surface pockets and catalytic cleft, we intend to investigate the impact of Heparin (Hep), Chondroitin sulphate (C4S) and PEG400 on FP2 activity. Our results show that Hep and C4S do not have any significant effect on FP2 activity for small substrates but PEG400 works as a potential inhibitor [Fig. S1(a)]. For the physiological substrate hemoglobin, both Hep and C4S reduces its proteolytic degradation by FP2 as observed in the SDS-PAGE based assay [Fig. S1(b)]. In contrast, PEG400 enhances rate of haemoglobin degradation by FP2 with almost complete elimination of intact Hb band under the same condition and time scale.

These multifaceted properties of PEG400 have encouraged us to probe its binding properties, mechanism of inhibition and regulation and modality of its interaction with the protease. We have performed the experiments with glycerol as control, to rule out non-specific solvent effects. No comparable effect was observed that confirm the specificity of PEG400 [Fig S1(c)].

#### Kinetic analyses

Inhibitory potency of PEG400 has been checked by measuring the activity of FP2 with increasing concentration of PEG400 using specific substrate D-VLK-pNA. The data was fitted to a single exponential binding model, yielding an IC_50_ value of 1.1% (v/v) (Table 1) [Fig. 1(b)].

**Table 1.**
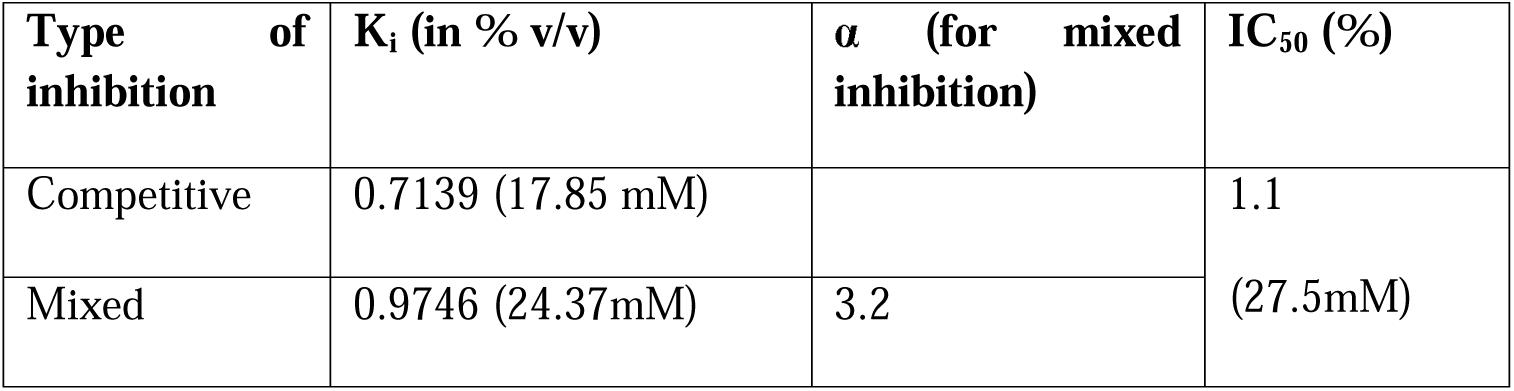
Inhibitor constants of PEG400.

Kinetic analysis using Michaelis–Menten [Fig. 1(c)] and Lineweaver Burk plots [Fig. 1(d)] show the concentration-driven aspect of inhibition of PEG400. At low concentration, the inhibition is almost competitive; however, at higher concentration, it propels a mixed-type inhibition with significant difference in V_max_ values. Dixon plot (1/V_0_ vs [I]) at a fixed concentration of substrate reveals a parabolic nature indicating that PEG400 has a multi-site co-operative interaction with FP2 [Fig. 1(e)]. The value of α, exceeding 1 (Table 1), is indicative of a mixed competitive type inhibition. The possible mechanism of this kind of kinetic behaviour is one where the inhibitor has higher affinity for the active site and occupies this site first; at higher concentration, it also tends to bind to one or more lower-affinity exosite/s. This additional binding at exosite(s) may induce some changes at the active site, altering its binding affinity associated with a co-operative behaviour of inhibition.

The intrinsic tryptophan fluorescence quenching of FP2 with increasing concentration of PEG400 yielded a K_D_ value of 1% (Fig S1d). The binding data are in agreement with the Ki value of PG400 (Table 1). Both the data are in mM range and corroborate low binding energy calculated from molecular mechanics study (Table 2).In order to ascertain whether the role of PEG400 as an inhibitor is specific to small peptide degradation or extends to other macromolecular substrates, we conducted an azocasein assay [Fig. S1e]. The results demonstrate a notable decrease in proteolytic activity in the presence of PEG400. Therefore, our findings indicate that PEG400 serves as an inhibitor of FP2 for both small and non-physiological macromolecular substrates.

**Table 2:**
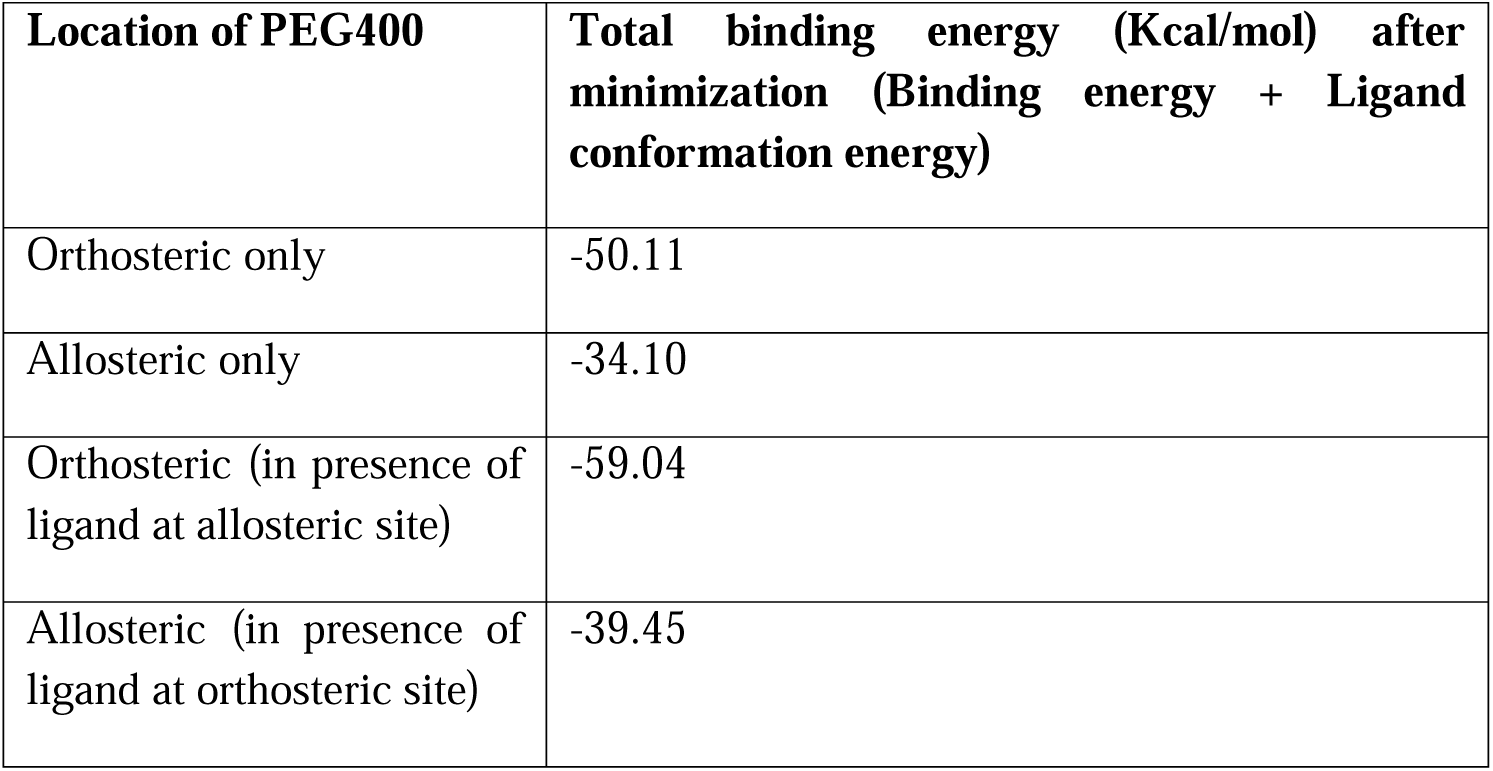
Binding energy of FP2 and PEG400.

| **Location of PEG400** | **Total binding energy (Kcal/mol) after minimization (Binding energy + Ligand conformation energy)** |
| --- | --- |
| Orthosteric only | -50.11 |
| Allosteric only | -34.10 |
| Orthosteric (in presence of ligand at allosteric site) | -59.04 |
| Allosteric (in presence of ligand at orthosteric site) | -39.45 |

To investigate the influence of PEG400 on other FP2-specific inhibitors, we also assessed the effect of PEG400 on two FP2-specific inhibitors, Leupeptin and human stefin-A (STFA). Each experiment was conducted by adding PEG400 after activation of FP2 and incubated briefly prior to adding respective inhibitors. In presence of PEG400, IC50 of Leupeptin changes [Fig. 1(f)] but that of STFA remained unchanged [Fig. 1(g)]. This suggests that Leupeptin being a potent inhibitor of cysteine proteases with higher binding affinity compared to PEG400, may replace PEG400 from the catalytic-site and probably positive allosteric effect of PEG400 due to binding at secondary site reduce the effective IC50 value of Leupeptin in presence of PEG400. In the case of specific proteinaceous inhibitor STFA, catalytic-site specific inhibitory loops and trunk of the inhibitor [9] may replace PEG400 from the catalytic cleft. In addition, the exosite interaction between FP2 and STFA may have a pivotal role in dislodging PEG400 from the catalytic cleft and may restrict the binding of PEG400 to the secondary site, incurring any allosteric effect.

#### Molecular Interactions and Binding Characteristics

##### Spectroscopic analyses

Mature FP2 structure contains four tryptophan residues and three of them are located in the vicinity of the catalytic cleft (Trp43, Trp206 and Trp210) while Trp24 is positioned at the back of FP2 [Fig. 2(a)]. In the crystal structure of FP2 [6,12], it was observed that the indole side-chain of Trp43, in spite of its sequence proximity with catalytic Cys42, is totally buried within the core of the L-domain while side-chains of the other three tryptophans are partially solvent-exposed [Fig. 2(a)]. Notably, Trp206 and Trp210 are facing towards the catalytic cleft whereas the cavity observed near the Trp24 residue is too small to accommodate a ligand like PEG400. Due to the proximity of Trp206 and Trp210 to the catalytic cleft, the fluorescence quenching is expected upon a ligand binding at the catalytic cleft due the change in microenvironment around these residues though these are not key residues of ligand/substrate/inhibitor interaction. The other two tryptophans are buried inside and unlikely to contribute. Therefore, Trp206 and Trp210 primarily account for the quenching signal and serve as a useful probes reflecting of PEG binding at the catalytic cleft.

**Figure 2.**
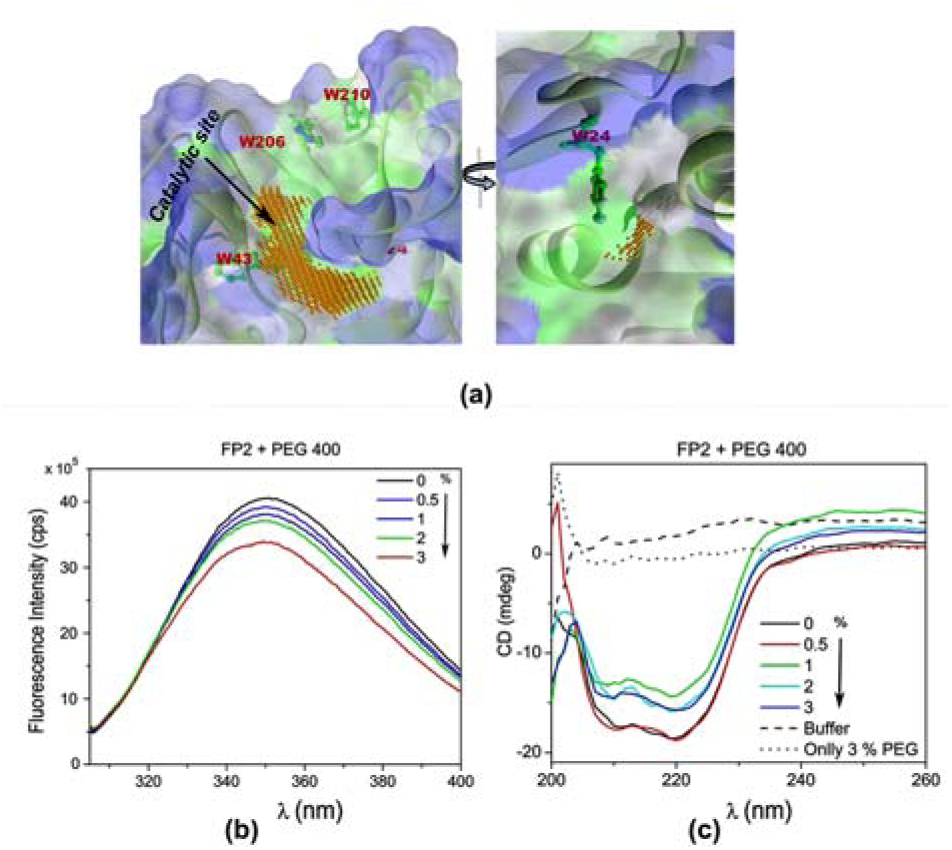
Spectroscopic analyses of FP2 and PEG400 interaction. (a) Tryptophan residue positions in FP2 structure. The nearby cavities are represented as orange dots (b) fluorescence and (c) Far-UV CD spectra of FP2 in presence of 0 - 3 % PEG400.

The emission maxima of tryptophan fluorescence of FP2 has been observed at 349.5 nm [Fig. 2(b)]. Titration of FP2 with PEG400 resulted in concentration-dependent reduction of the fluorescence intensity along with minimal red shifts of the emission maximum [Fig. 2(b)]. The indole side-chain moiety of the tryptophan residue is very sensitive to changes in surrounding micro-environmental conditions and thus is indicative of changes in protein conformation upon ligand binding. The observed fluorescence quenching with red-shift of absorption maxima (349.5→351.5 nm), confirm shifting of trytophan residue(s) to a more solvent exposed position due to binding of PEG400. Since two Trp residues are in the proximity of the catalytic cleft of FP2, the catalytic site would be a probable binding site of PEG400, which corroborate the enzyme inhibition kinetic study of FP2 in the presence of PEG400.Far-UV circular dichroism (CD) spectra reveal a PEG400 concentration-dependent change in the CD signals of FP2, indicating changes in the secondary structure without causing global unfolding of the protein [Fig. 2(c)]. However the trend in spectral change is not uniform with increase in PEG400 concentration. A significant reduction of CD signal is observed for 1% PEG inferring partial loss of secondary structure of FP2. However, the CD signal shows partial recovery of secondary structural elements when concentration of PEG400 was further increased to 2-3%.

#### Hypothetical mechanism of inhibitor binding

Enzyme kinetic analysis reveals that at low concentration, PEG400 acts as a competitive inhibitor confirming its binding at the FP2 catalytic cleft and this is in agreement with fluorescence spectroscopy observations. However, kinetic analyses further reveal the possibility of presence of multiple binding sites of PEG400 at higher concentration accounting for a mixed-type of inhibition pattern. Molecular docking and cavity analyses have been performed to gain structural insight into PEG400 binding, using the crystal structure of FP2 previously determined by our group [6]. Among the first top ten docking poses determined by Autodock vina [31] eight poses are located at the orthosteric site and two at an allosteric site within the FP2 L-domain [Fig 3(a), 3(b)]. PEG400 tends to bind in extended as well as in folded conformations [Fig 3(b)].

**Figure 3.**
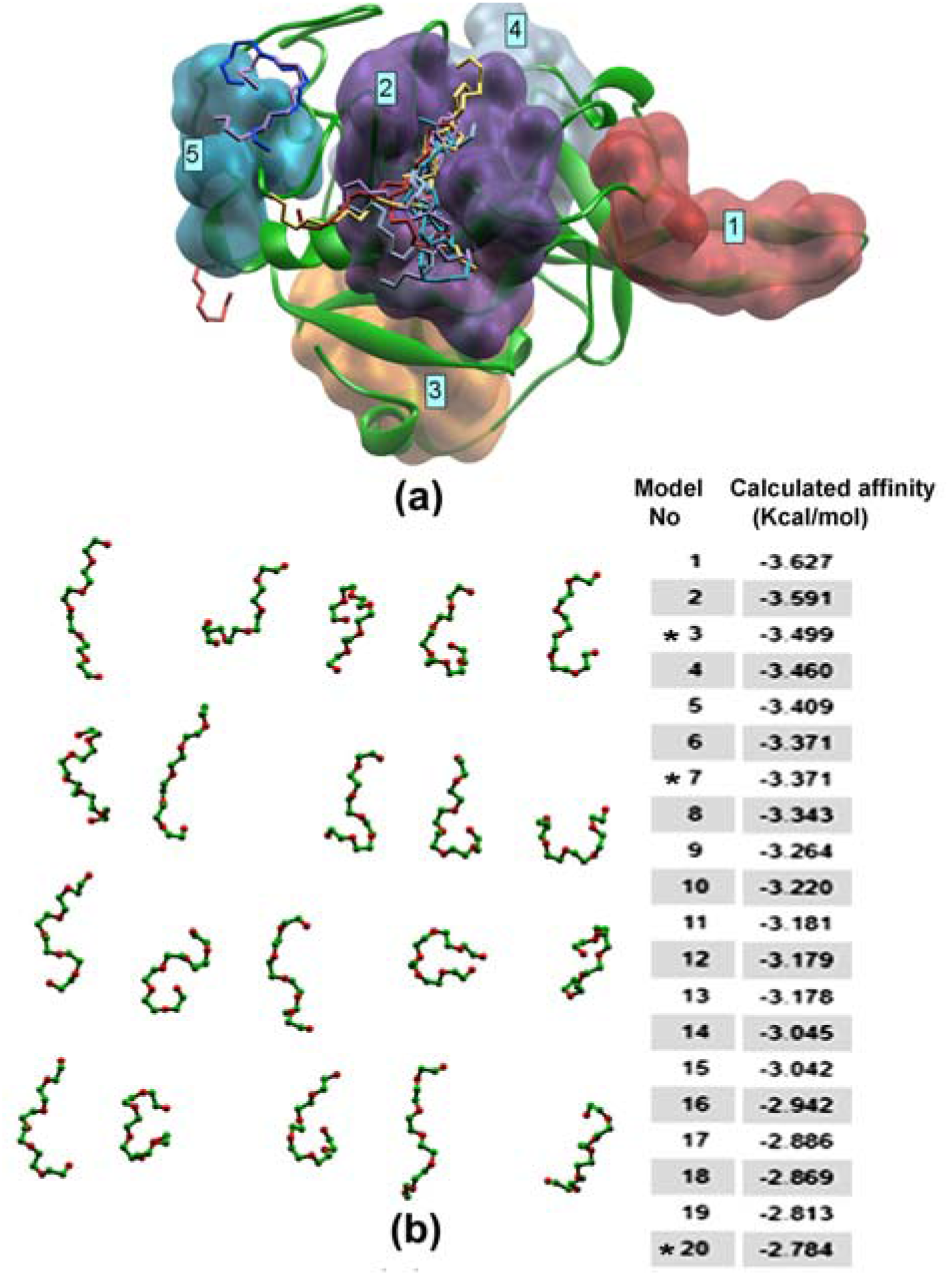
Docking of PEG400 and cavity analysis of FP2. (a) Structural representation of FP2 showing the five cavities identified by CavityPlus, marked according to their calculated cavity volume. Among these, cavity number 2 corresponds to the orthosteric (catalytic) site of FP2. Docked PEG400 molecules are shown in stick model. (b) Summary of molecular docking results displaying the top 20 binding poses of PEG400, ranked by docking score as shown in the embedded table. Poses at predicted allosteric sites are marked with an asterisk (*), indicating potential alternative regulatory binding region outside the catalytic cleft.

CavityPlus web server (http://pkumdl.cn:8000/cavityplus/<u>#/</u>) has been used subsequently to identify probable allosteric sites in FP2. Fig. 3(a) illustrates five predicted ligand binding pockets in FP2. Among these, site 2 is the orthosteric site, while site 5 has been identified as an allosteric site in the L-domain of FP2 where PEG400 has two docking poses. Subsequently, with site 2 set as the orthosteric site, the allosteric potential of the remaining four sites were predicted with the CorrSite submodule of CavityPlus. It shows the highest score of 1.5 for site 5, significantly higher than the next value of 0.86 for site 1. Based on these combined studies, the most probable allosteric site for PEG400 binding has been identified as site 5 which is located in close proximity of the orthosteric site and separated by two surface loops [Fig. 3(a)]. The same site is also identified in top-scoring allosteric pocket using Allosite tool [32]. This further validates the predicted allosteric binding pocket for PEG400.

Docking studies prompted us to select a ligand conformation with the highest score for both orthosteric (pose 1) and allosteric sites (pose 3) [Fig. 3(b)] to perform molecular mechanics calculations. Individual FP2-PEG400 complexes, as well as the FP2 complex containing two PEG400 molecules at orthosteric and allosteric sites were energy minimized and their binding energies were subsequently calculated. The minimized structures reveal that PEG400 at the orthosteric site has higher binding affinity compared to the allosteric site (Table 2). Comparing the binding energy and cleft volume analyses, it was observed that PEG400 forms a more stable mode of interaction with the FP2 catalytic cleft compared to the shallow allosteric pocket [Fig. 4(a) and (b)].

**Figure 4.**
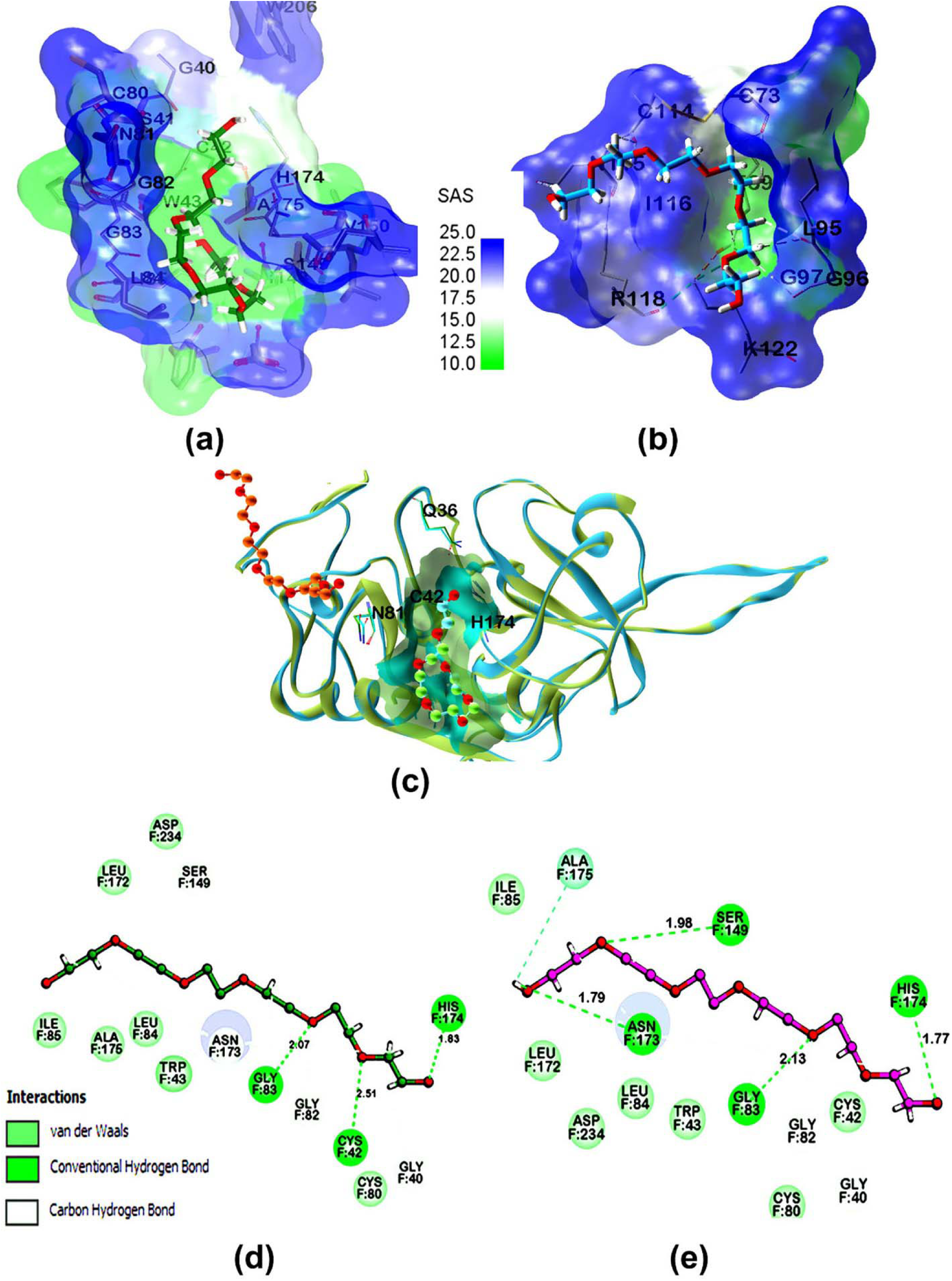
Interaction of top scoring docking poses of PEG400 molecules after energy minimization at. (a) orthosteric and (b) allosteric sites with solvent accessible surface of FP2. (c) Superposition of catalytic and allosteric site in absence and presence of PEG400 bound at allosteric site. Orthosteric Volume reduced 284.1Å^3^ (light green) →268.5 Å^3^ (sky blue) upon PEG binding at allosteric site. (d) 2D interaction diagram of PEG400 with FP2 comparing the binding environment in absence and (e) presence of PEG400 at allosteric site.

This study further reveals that there is a change of binding pattern between PEG400 and FP2 when both the sites are occupied [Fig. 4 (c), (d) and (e)]. Due to PEG400 binding at allosteric site, a conformational change is observed in the loop between G79 and N81, located at the interface of orthosteric and allosteric sites. This change leads to a better packing of PEG400 at the catalytic cleft by lowering of interacting cavity area [Fig. 4 (c)]. A minute alteration of residues lining the catalytic cleft is thus induced, stabilizing both the ends of PEG400 through conventional hydrogen bonds [Fig. 4 (d) and (e)] which further lowers the interaction energy (Table 2). Additionally, binding of PEG400 at allosteric site induces a side-chain movement of Gln36, an important residue involved in aligning H174 side-chain to form Cys42^-^-His174^+^ catalytic dyad, is observed [Fig 4 (c)]. These data corroborate with our kinetic study where a multi-site interaction of PEG400 is expected to be co-operative.

We further performed docking study of Leupeptin and FP2 to correlate the kinetic data. The study depicts that leucine residue (P2) of the inhibitor occupies the specificity determining S2 subsite of FP2 [Fig S2a]. Comparing the mode of interaction of PEG400 and FP2 with that of Leupeptin and FP2 [Fig S2b] shows PEG400 only occupies P1, P2 and P3/P4 region while Leupeptin has additional interaction with FP2 catalytic cleft covering its Arginyl and second Leucyl moieties. These additional interactions contributed higher binding energy (-69.06 Kcal/mol compared to -50.11 Kcal/mol for PEG400) and affinity of Leupeptin to the enzyme. This study confers that Leupeptin may replace PEG400 (which was incubated with FP2 prior to adding Leupeptin) and allosteric effect due to binding of PEG400 at allosteric pocket may influence lower IC50 value of Leupeptin in presence of PEG400.

PEG molecule typically binds to hydrophilic, non-specific shallow surface grooves or solvent channels in the protein. However, in our earlier crystal structure of FP2 [6], we observed PEG molecule at the catalytic cleft. This indicates that PEG can engage not only ‘soft’ nonspecific interaction but also in more defined, site-specific interaction with proteins. Due to its structural flexibility, it can reach to a solvent exposed well-defined cleft of a protein with structural adoptability according to the cleft shape if the chain length matches [33] as it is observed in this study. If the binding of PEG400 at the allosteric site of FP2 is considered as soft interaction, then higher binding affinity for the catalytic cleft could be considered as specific interaction.

The additional work on comparative analysis with leupeptin further supports this interaction, as PEG400 overlaps with leupeptin binding at specificity-determining S2 subsite as well as S1 subsite of the catalytic cleft.

#### Molecular Dynamics simulation

We performed a short MD simulation to understand the dynamic trends and correlated motion of allosteric and active site residues. Production run of 100 ns has been executed for FP2, with coordinates saved every 20 ps (total 5000 frames) for subsequent RMSD, RMSF, radius of gyration (Rg), hydrogen-bond and principal-component analysis. The simulation trajectory reveals a stable potential energy (Fig S3a). The average Cα RMSD has been calculated against the trajectory midpoint conformation (at 50 ns) as reference structure (Fig S3b). The RMSD data shows higher values for the 1st 10 ns trajectory, fluctuation upto 60 ns while rest of the trajectory beyond 60 ns has reasonable lower values in the range of 1.5 Å. The root mean square fluctuation (RMSF) plot (Fig S3c) primarily shows less flexible L-domain compared to R-domain. Significant high fluctuations are observed for the nose region and Hb-binding loop/region (Fig S3c). The enzyme retains its overall compactness throughout the trajectory as depicted from radius of gyration plot (Fig S3d) with Rg value ranges between 18.06-18.13 Å. Number of H-bonds are also consistent (Fig S3e) in the entire trajectory.

#### Principal component analysis (PCA)

PCA analysis was performed using the entire 100 ns production time of the MD trajectory to identify the dominant motions and the transitions between conformations in the course of the simulation of FP2. This approach allows to describe the essential dynamics motions of the molecule, where low-frequency eigenvectors with relatively higher eigenvalues, explaining most of the movements and the calculated parameter principal components PCs are ranked accordingly. Figure 5a represent a 3D plot of first three principal components PC1 (63.2%), PC2 (18.5%) and PC3 (10.8%) respectively, those cover most of the variation. The observed conformations can be grouped into three clusters with population density highest for cluster 1 followed by cluster 2 and cluster 3 (Fig 5a). The most densely populated cluster 1 contains the conformations for the last half of the trajectory, while clusters 2 and 3 are from the first half of the trajectory. A constant drift is observed from one group of conformations to another group with smooth transition from cluster 3→2→1. The cluster analysis of the trajectory is in good agreement with rmsd values in the trajectory where last half of the trajectory have stable RMSD value (Fig Sb) containing the conformations of cluster 1. Superposition of representative structures (Fig 5b), selected from the central region of each clusters, shows overall Cα rmsd value in the range of 3-3.5 Å with dominating differences in the Hb-binding loop/region of R-domain and in the nose region (Fig S3f) as observed in the sequence alignment based on the structural superposition. We have evaluated the allosteric binding site by docking PEG400 and the topmost poses from each structure of the clusters have been compared (Fig 5c). The study reveals that in all the three structural representatives from the clusters, the interacting regions of PEG400 have overlapping with the allosteric site identified in FP2 crystal structure (Fig 5c). However the Autodockvina score is highest for cluster 1 and comparable with FP2 crystal structure with similar PEG400 interaction pattern (Table S1). This study further recommend the allosteric site for PEG400 since cluster 1 contains the highest number of population in the MD trajectory. The PCA analysis of the protein conformational space reveals the direction of the observed movements. Figure 5d shows which parts of an FP2 molecule contribute most to a PC1, PC2 and PC3 i.e., the highest-amplitude motion modes. The three PCs involve motions, as indicated by square fluctuation, in several regions of a protein including the Hb-binding loop/region, nose regions and other protein parts labelled in Figure 5d. Directions of their oscillation are visualized on a porcupine plot (Fig 5d) by double-headed arrows. The predominant character of conformational changes in essential dynamics are a rocking motion nose region and hinge-bending motion of Hb-binding loop causing the shift of the concerned back-bone in the observed directions.

**Figure 5.**
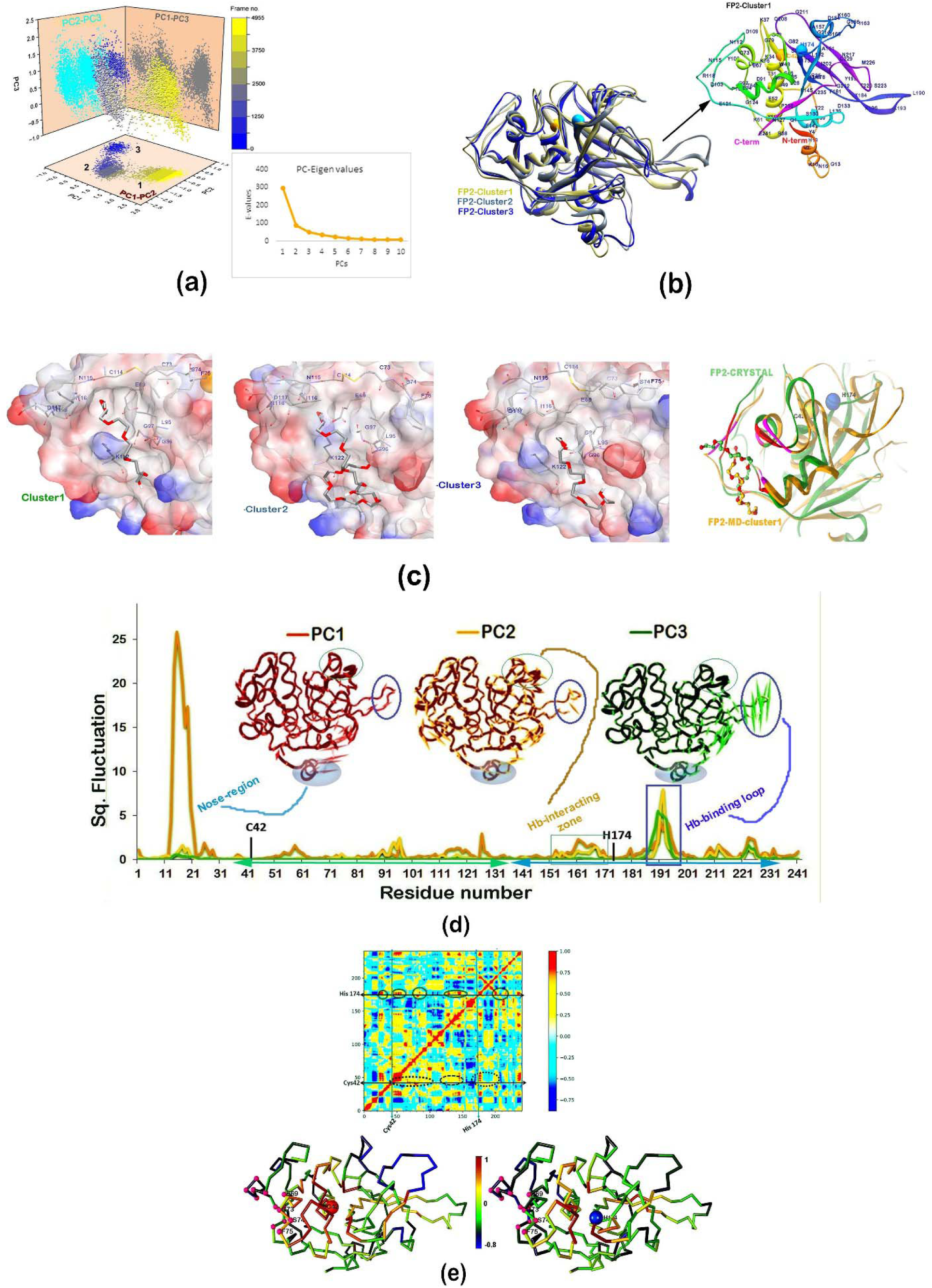
Principal component analysis. a) 3D plot of PCs 1, 2 and 3(left panel) with the colour scheme corresponding to frame number of the trajectory; from the beginning (blue) to the final 5000 frame (yellow). The conformations are clusters in three segments and are marked as cluster 1, 2 and 3 according to population density. The plot (right panel) shows variation of top ten PCs captures from the data, b) superposition of three representative structures picked up from the central region of three clusters. The representative structure from cluster 1 has been shown independently with residue_id marked intermittently (right panel) to understand the region in the structure. The structure has been presented as ribbon diagram with rainbow colour scheme, c) docking analysis of PEG400 (top score position) with three conformations of FP2 clusters. The right panel shows superposition of cluster1 structure with FP2 crystal structure, d) square fluctuation of residues of FP2 along PC1, PC2 and PC3 with colour coding orange, yellow and green respectively. Porcupine plots are embedded to visualize the direction of PC1, PC2 and PC3 obtained from the PCA. The double-headed arrow direction in each Cα atom represents the direction of motion, while the length of arrow (in relative scale for each structure) characterizes the strength of the associated movement, (e) inter-residue motion illustrated in cross correlation map of the Cα atoms of FP2. Positive values (red spectrum) indicate correlated motion; negative values (blue spectrum) indicate anti-correlated motions amongst residues. Cross-sections at C42 and H174, the catalytic dyad residues, are marked. Lower panel shows ribbon diagram of FP2 crystal structure with colour coding according to C42 and H174 cross-sections. The respective C42 and H174 Cα s are represented as sphere and marked. To understand correlation between dyad residues and allosteric site, the allosteric site residues are represented as smaller spheres.

#### Correlation of allosteric site with catalytic site

While the analysis of RMSF identifies the most flexible parts of a molecule in general, it does not reveal if these parts move in concert or independently, and it cannot distinguish between directed or random movements. The dynamic cross-correlation map (calculated for PCA) would reveal the correlated dynamics of structural fluctuations, and in particular, provide the key to identify the correlation between distal regions of a protein. In this map, red indicates positive correlations, while blue indicates negative correlation. A cross sections at Cys42 and His174, the catalytic dyad residues are shown in the upper panel of Figure 5f and the corresponding ribbon diagrams were presented with colour red to blue for positive and negative correlation normalised as 1 to -1. The correlated motion between catalytic Cys42 and allosteric residues has been observed in the cross correlation map of FP2, (Fig 5e). On the other hand cross correlation between H174 and allosteric residues are insignificant. The study proposed that the local structural/dynamic changes at the allosteric site upon binding with a ligand induce changes at catalytic Cys42 residue as they are positively correlated.

#### PEG400 promotes Hemoglobinase activity of FP2

##### SDS-PAGE based assay

Host hemoglobin (Hb) is the primary substrate of FP2 in parasite life cycle during trophozoite stage. Our analysis shows that 5.1 µM FP2 is sufficient to almost completely degrade 1.1 µM tetrameric Hb within 30 min under the optimized condition, highlighting the catalytic efficiency of FP2 [Fig. 6(a)]. We assessed the hemoglobinolytic activity of FP2 in the presence of 0.5% to 5% PEG400. The study indicates PEG400 enhances hemoglobinolytic activity of FP2 [Fig. 6(b)]. The corresponding band intensities for the intact Hb at 14 kDa have also been estimated. The faster disappearance of Hb bands around 14 kDa with increasing concentration of PEG400 signifies that PEG400 acts as an inducer of hemoglobinase activity of FP2 and this observation is contrary to its inhibitory behavior for a smaller substrate of FP2. We have performed SDS-PAGE based time-course profiling of hemoglobin degradation by FP2, both in presence and absence of PEG400. The results revealed an accelerated breakdown of Hb in the presence of PEG400. The band intensity analysis of intact Hb was quantified and the results are presented as bar diagrams for clearer comparison [Fig. 6(c)].

**Figure 6.**
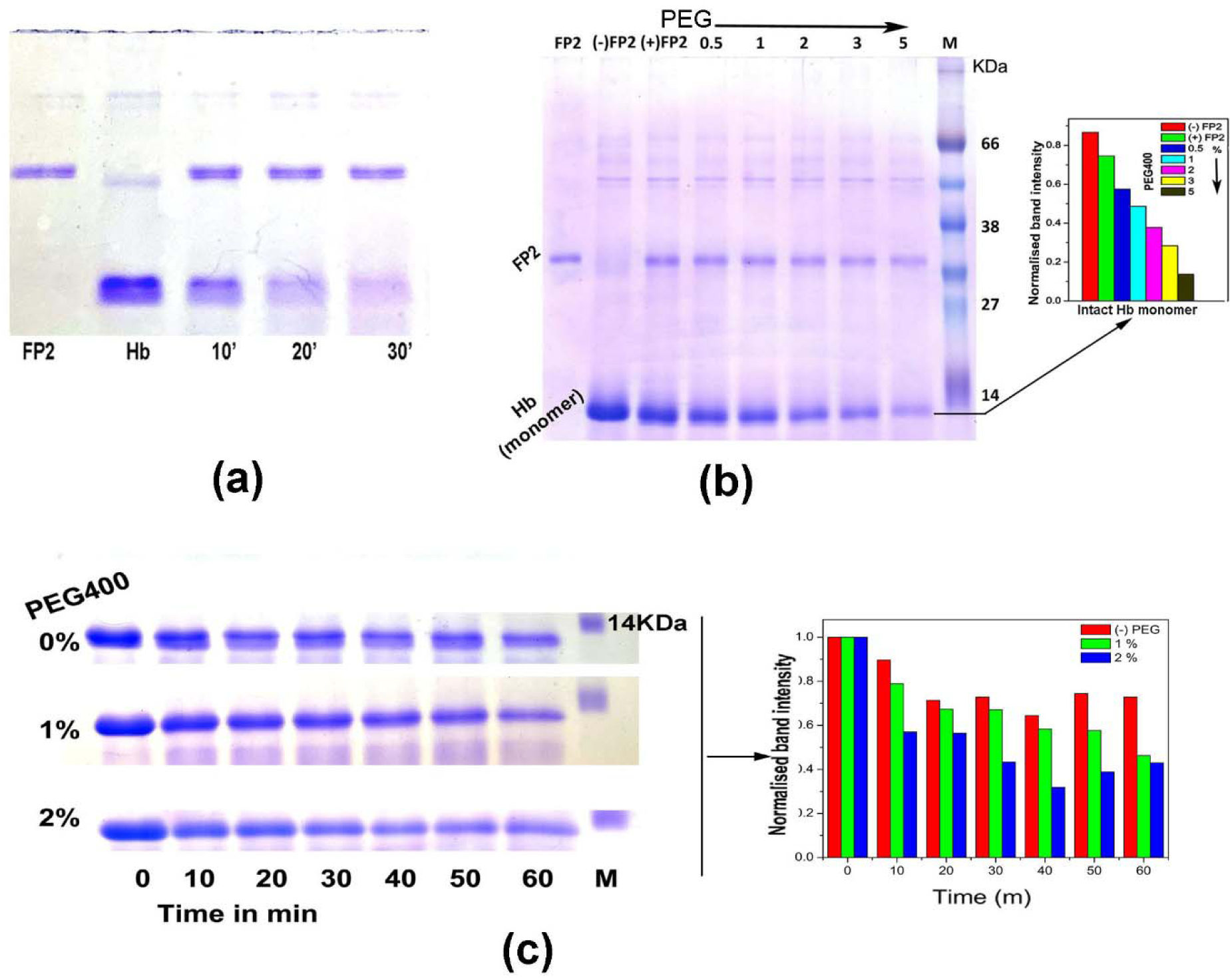
Haemoglobin degradation analysis. (a) 5.1 µM FP2 degraded 1.1 µM tetrameric Hb within 30 m. (b) SDS-PAGE analysis after 15 min of incubation at 37°C where FP2, (-) FP2, (+) FP2 and 0.5 - 5 denotes FP2 only, Hb only, Hb with FP2 and Hb with FP2 in presence of 0.5 - 5 % PEG400. (c) SDS-PAGE of Hb (7.1 µM) incubated with FP2 (3.6µM) at 37°C for 0-60 m in presence of 0-2% PEG400. Normalised band intensity of intact monomeric Hb are shown in the right panel of (b) and (c).

##### Spectroscopic analysis

Tryptophan fluorescence spectroscopy has been used to understand the binding of PEG400 with Hb alone and in complex with FP2 as Hb contains six tryptophan residues in its tetrameric structure while FP2 contains four. The results shows spectral quenching of Hb fluorescence with minimal red-shift on emission maxima [Fig 7(a); Table S2] with increasing concentration of PEG400. This suggests a realignment of Trp residues buried in hydrophobic cavities to a more hydrophilic environment, confirming binding of PEG400 to Hb. In contrast, a blue-shift and increased fluorescence intensity has been observed for Hb-FP2 complex for 0.5% PEG400 [Fig 7(a)]. Such shifts of fluorescence peaks, known to result from the tryptophan(s) becoming more buried and shielded away from the hydrophilic environment, are indication of changes of the local environment around the Trp residue(s) either in Hb or in FP2 or in both upon mutual interaction. Notably, neither of the individual proteins shows such observation [Fig 2(b) and 7(a)]. However when concentration of PEG400 was further increased to 1% and 2%, the spectra is almost super-imposable with that of the complex without PEG400 and further quenching is observed at 3%. Since there are ten Trp residues (6+4) in the Hb-FP2 complex, the complexity of the fluorescence spectral phenomena restricts from having any conclusive opinion alone from this study. Therefore we performed circular dichrism (CD) and absorption spectroscopy in the Soret region for further investigating the mode of binding.

**Figure 7.**
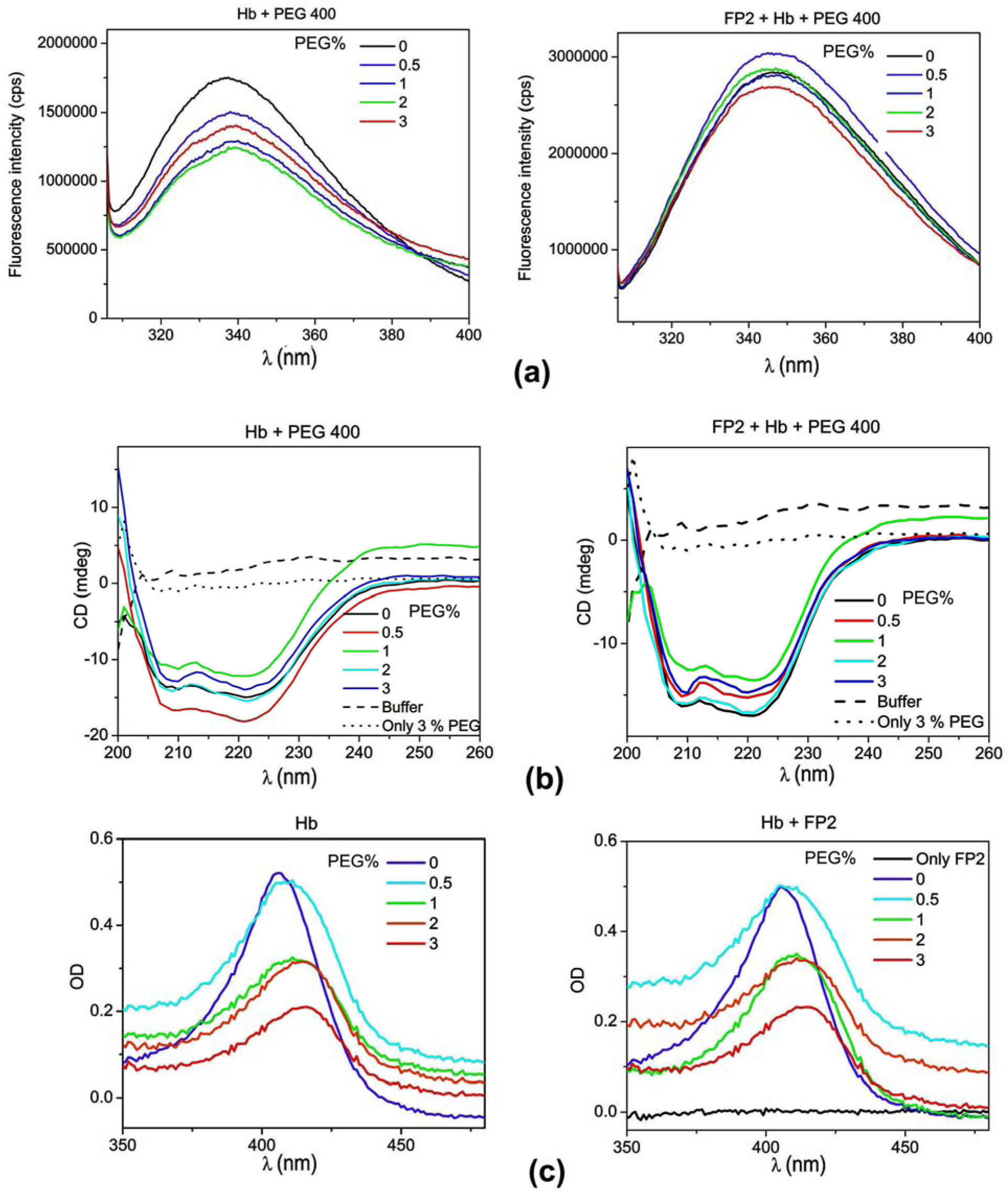
Spectroscopic analyses of Hb and FP2-Hb complex. (a) Tryptophan fluorescence in the presence of 0 - 3 % PEG400 for Hb and mixture of Hb and FP2. (b) Far-UV CD spectra of Hb and 1:1 mixture of FP2 and Hb, recorded at pH 8.0 and 25° C in the presence of 0 - 3 % PEG400. (c) Near-UV and Soret absorption spectra of Hb in presence of 0 - 3 % PEG400 (denoted as 0, 0.5, 1, 2, 3) and Hb-FP2 mixture, under the same PEG400 concentration, recorded at pH 8.0 in Tris buffer.

Far-UV CD spectra of Hb shows an interesting observation; at 0.5% PEG400, the CD signal at wavelength characteristics for α-helix (208 and 222 nm) increased significantly and then decreased to nearly the original level at 1%. These dramatic changes signify that secondary structural changes of Hb attenuate in reverse direction at 0.5% and 1% PEG400 [Fig 7(b)]. A similar reduction of α-helical content is observed in Hb-FP2 complex at 1% PEG400 [Fig 7(b)] and as well as FP2 alone [Fig 2(c)][Fig S4]. Therefore 1% PEG400 is a critical concentration where individual FP2, Hb and their complex significantly lost their secondary structure [Fig S4]. Interestingly, original structures are partially restored when the concentration of PEG400 is further increased [Fig 2(c), 7(b)].

We demonstrated the impact of PEG400 on the absorption band (Soret region) of Hb [Fig. 7(c)], as well as on a combination of Hb and FP2 [Fig. 7(c)]. With increase in concentration of PEG400, a reduction in intensity of soret peak and a red shift in λ_max_ is observed in both cases of Hb and the mixture of Hb and FP2 [Table S3]. These spectral changes suggest dislocation of heme group from the respective Hb binding pocket with increasing PEG400 concentration. A sharp reduction of Soret peak intensity has been observed at 1% PEG400 similar to CD spectra.

To comprehend the dynamics of the changes in the heme group of Hb in the presence of PEG400, we monitored the absorbance at 406 nm over time in the presence of 0.5 - 3% PEG400. The rate constant shows biphasic kinetics of heme delocalisation, a faster rate in the first 2 minutes and a slower rate beyond 2 minutes [Fig. S5 and Table S4]. These findings suggest a shift in the reaction mechanism or the stability of intermediate states over time. The initial rapid phase (0–2 minutes) shows a sharp increase of rate constant k_1_ to a maximum value at 1% PEG400 [Table S4], indicating a fast conformational change in this concentration. In contrast, the slower second phase (after 2 minutes) [Table S4], exhibits a concentration-dependent increase of rate constant k_O_, pointing to a gradual, PEG-induced structural reorganization. This biphasic characteristic may be attributed by to two types of globin chains α and β in hemoglobin.Overall, these findings suggest that PEG400 does not induce total denaturation of Hb or Hb-FP2 complex. Instead, it induces conformational changes in both FP2 and Hb that alter the environment of the heme group. These conformational modifications may be the underlying reason for the enhanced rate of catalysis of FP2 against Hb in presence of PEG400.

##### Understanding the enhanced hemoglobinase activity of FP2 in presence of PEG400 at molecular level

We assessed the mode of interaction between PEG400 and both with Hb and Hb-FP2 complex through docking studies. Crystal structure of Hb (pdb_id: 2DN1) and a model of Hb-FP2 complex reported earlier in our study [6] were used for docking with PEG400. The study identified PEG400 binding site at the central cavity and near the heme-binding region [Fig 8(a)]. These interactions suggest that PEG400 binding may induce structural changes in the globin part as well as heme location of Hb, corroborating with the spectroscopic studies. In addition to these sites of Hb, PEG400 molecules were found at the interface of Hb-FP2 complex with additional interactions [Fig 8(b)] bridging the two molecules and creating a better stabilizing effect in the complex.

**Figure 8.**
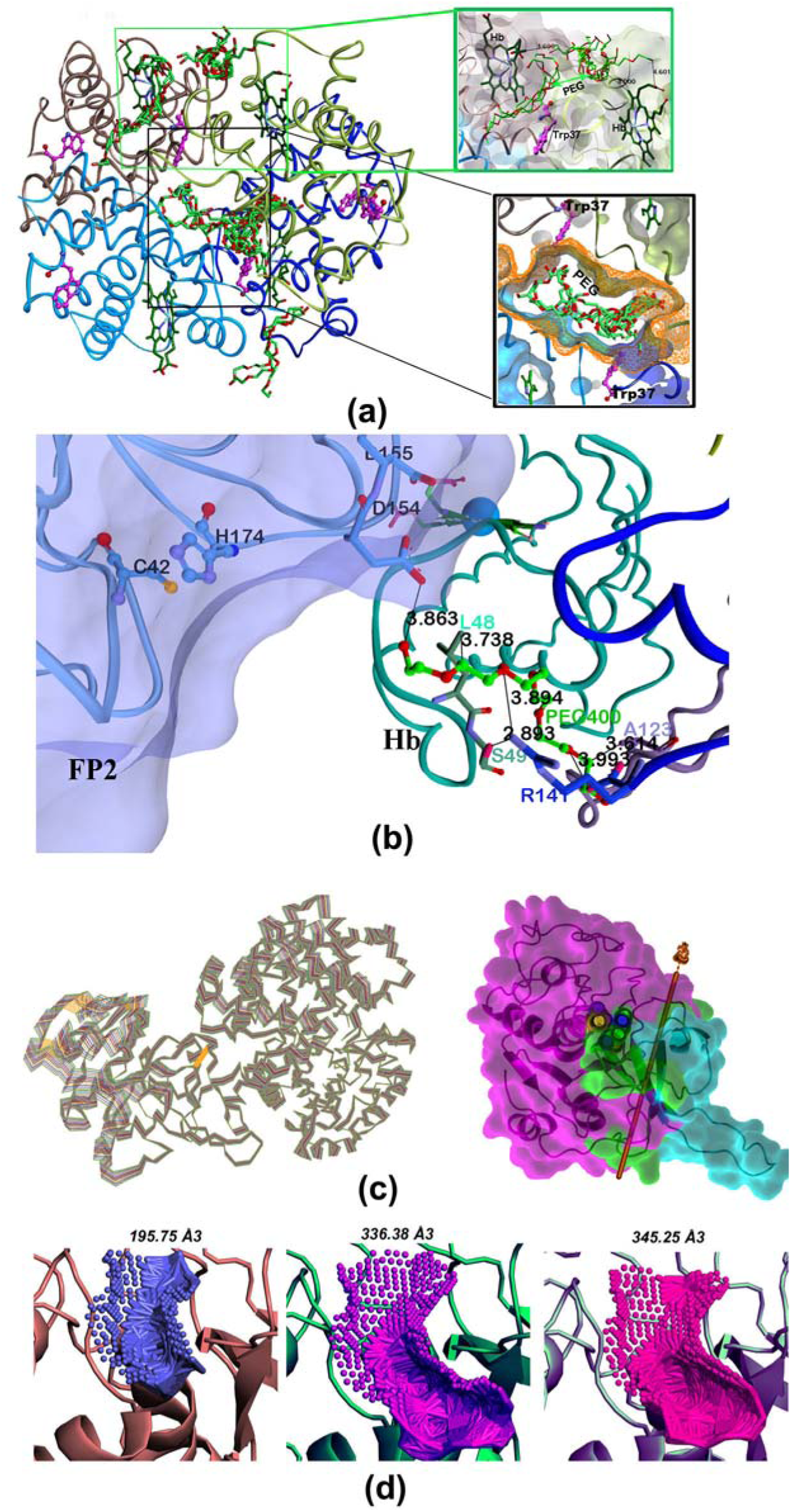
Docking of PEG400 in Hb and FP2-Hb complex and normal mode analysis, (a) Docking analysis showing the predicted PEG400 binding site within the central cavity of Hb and (b) located in close proximity to the heme-binding region. (c) NMA of FP2-Hb complex illustrating the dominant collective motions that may facilitates substrate binding and product release. Right panel shows a hinge-bending motion within FP2. The mobile domain, fixed domain and hinge-bending region are colored as magenta, sky blue and green respectively with the axis of rotation is shown in the figure (d) Structural visualization of the catalytic cleft demonstrating its reduction and expansion of catalytic cavity as a consequence hinge-bending residues.

To understand the effect of PEG400 on the dynamics of Hb-FP2 binding, we performed normal mode analysis (NMA) of the Hb-FP2 complex, in presence and absence of PEG400 [Figs 8(c)]. The study reveals that FP2 shows a hinge-bending motion in a similar fashion in both cases. During this motion, entire L-domain and a part of R-domain (moving domain, magenta) rotate around an axis passing through the region (hinge bending part; green) from where Hb-binding loop protrudes (fixed part; sky blue) [Fig 8(c)]. The Hb-binding loop acts as an anchor, tethering FP2 to Hb. This hinge-bending motion probably enables the FP2 catalytic centre to come in proximity to bound Hb for catalysis to occur. Some residues in the moving domain are situated on the inner-wall of catalytic cleft at the R-domain side which also includes catalytic dyad residue His174. The reduction and expansion of catalytic cavity is observed due to the presence of bending residues which ranges from the minimum volume of 195.75 Å^3^ to the maximum of 345.25 Å^3^ with the crystal structure occupying an intermediate of 336.38 Å^3^ [Fig. 8(d)]. The squeezing of the catalytic cleft with reduced volume from 336.38 Å^3^ to 195.75 Å^3^ during the hinge bending motion upon Hb binding, in all probability, leads to displacement of PEG400 from the catalytic cavity since PEG400 docked catalytic cleft of FP2 has a minimum volume of 268.5 Å^3^ [Fig 4(c)] and thus enable FP2 to become proteolytically active to degrade bound Hb. This is a plausible explanation for PEG400 not inhibiting hemoglobin degradation even upon prior incubation of FP2 with PEG400. The L-domain, on the other hand, undergoes a movement as a rigid part during the hinge-bending motion and therefore dislodging PEG400 from the allosteric site is unlikely. As a result PEG400 mediated allosteric regulation may continue to influence Hb proteolysis. We also observed the interaction energy, calculated for FP2 and Hb complex in presence and absence of PEG400, are comparable [Fig S6]. Therefore, observed enhanced hemoglobinase activity of FP2 in presence of PEG400 may be attributed by one or a combination of reasons: a) NMA results showing the presence of PEG400 restricting the hinge bending motion (rotation is reduced from 9° to 7°) [Fig. S7] as a whole and PEG400 molecules at the interface of FP2-Hb interlacing two molecules, stabilizing the Michaelis complex thereby enhancing the rate of catalysis, b) PEG400 induced conformational changes in Hb, as observed in the combined spectroscopic studies, may be conducive for easy catalysis, c) persistent binding of PEG400 at the allosteric site may positively regulate Hb catalysis by maintaining a catalytically favorable conformation of FP2.

#### Effect of high molecular weight PEG

To evaluate if molecular size of PEG have the same influences in modulation of FP2 activity, we examined higher molecular weight PEGs of 4000 and 6000 and compared them with PEG400. Hydrolysis of the D-VLK-pNA substrate [Fig. 9(a)] showed that PEG4000 did not induce inhibition, while PEG6000 has inhibitory effect although weaker than that of PEG400 [Fig. 9(a)]. PEG6000 acted as an activator during Hb degradation [Fig. 9(b)], whereas PEG4000 had no observable effect. For azocasein substrate [Fig. S1e], neither PEG4000 nor PEG6000 caused inhibition. Thus, PEG4000 shows no effect across substrates whereas PEG6000 shows effect similar to PEG400, but to a lesser extent. In summary, effect of PEG 6000 is similar to PEG400. It acts as inhibitor for small peptidyl and non-specific macromolecular substrates (e.g., azo-caesin) but activates hemoglobin catalysis.

**Figure 9.**
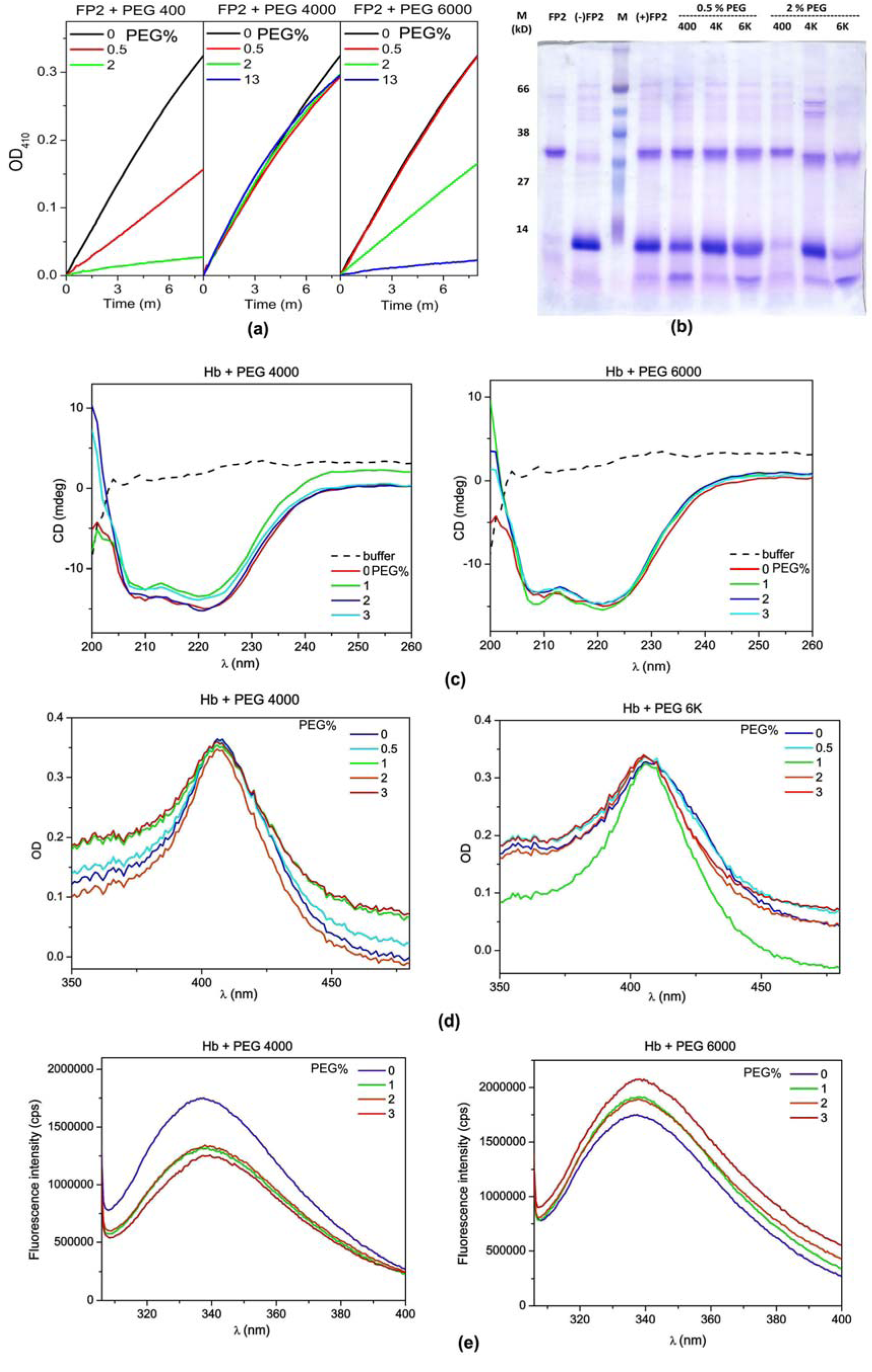
Effect of high molecular weight PEG. (a) OD at 410 nm against time using FP2 and the chromogenic substrate D-VLK-pNA in the presence of PEG400, PEG 4000 and PEG 6000. (b) Hb degradation analysis after incubation at 37°C for 25 min in the presence of different PEGs using 15% SDS PAGE. (c) Far-UV CD spectra of Hb, recorded at 25°C in the presence of 0 - 3 % (denoted as 0, 0.5, 1, 2, 3) PEG 4000 and PEG 6000. (d) Near-UV and Soret absorption spectra of Hb in presence of 0 - 3 % PEG 4000 and PEG 6000. (e) Tryptophan fluorescence for Hb in the presence of 0 - 3 % PEG 4000 and PEG 6000.

Far-UV CD and absorption spectra at Soret region [Fig, 9(c), 9(d)] shows minimum change in secondary structure and heme orientation in presence of high molecular weight PEGs. Intrinsic Trp fluorescence spectra shows a sharp quenching in the presence of PEG4000 [Fig 9(e)] indicating exposure of Trp residue(s) to a polar environment, On the contrary, fluorescence intensity increases in presence of PEG6000, may be owing to prevention of Trp-heme energy transfer and making a more hydrophobic environment to the Trp residue(s). The spectroscopic studies as a whole reveal nominal structural dynamics of Hb in presence of higher molecular weight PEGs, however local unravelling of the structure around the Trp side-chain or change of rotamer position could not be ignored. A more detailed investigation of these interactions is omitted here for brevity.

### *In silico* mutational analysis

The identified allosteric site for PEG400 appears to be amphipathic nature with a disulphide bond contributed by residues C73 and C114 at the base of the cleft and wall is hydrophobic at one side and hydrophilic at the other (Fig 10a). The central part of the ligand is positioned at the core of the allosteric cleft while the flanking end of PEG400 is solvent exposed (Fig 10a). PEG400 at allosteric site is stabilized by hydrophobic and favourable electrostatic interactions (Fig 10b). The disulphide bond at the base (Fig 10a) is highly conserved in the C1 family as depicted from amino acid sequence alignment with human cathepsins L, K, S and B which are abundantly expressed in almost all human cells (Fig 10c). Rest of the residues padding the cleft are mostly non-conserved.

**Figure 10:**
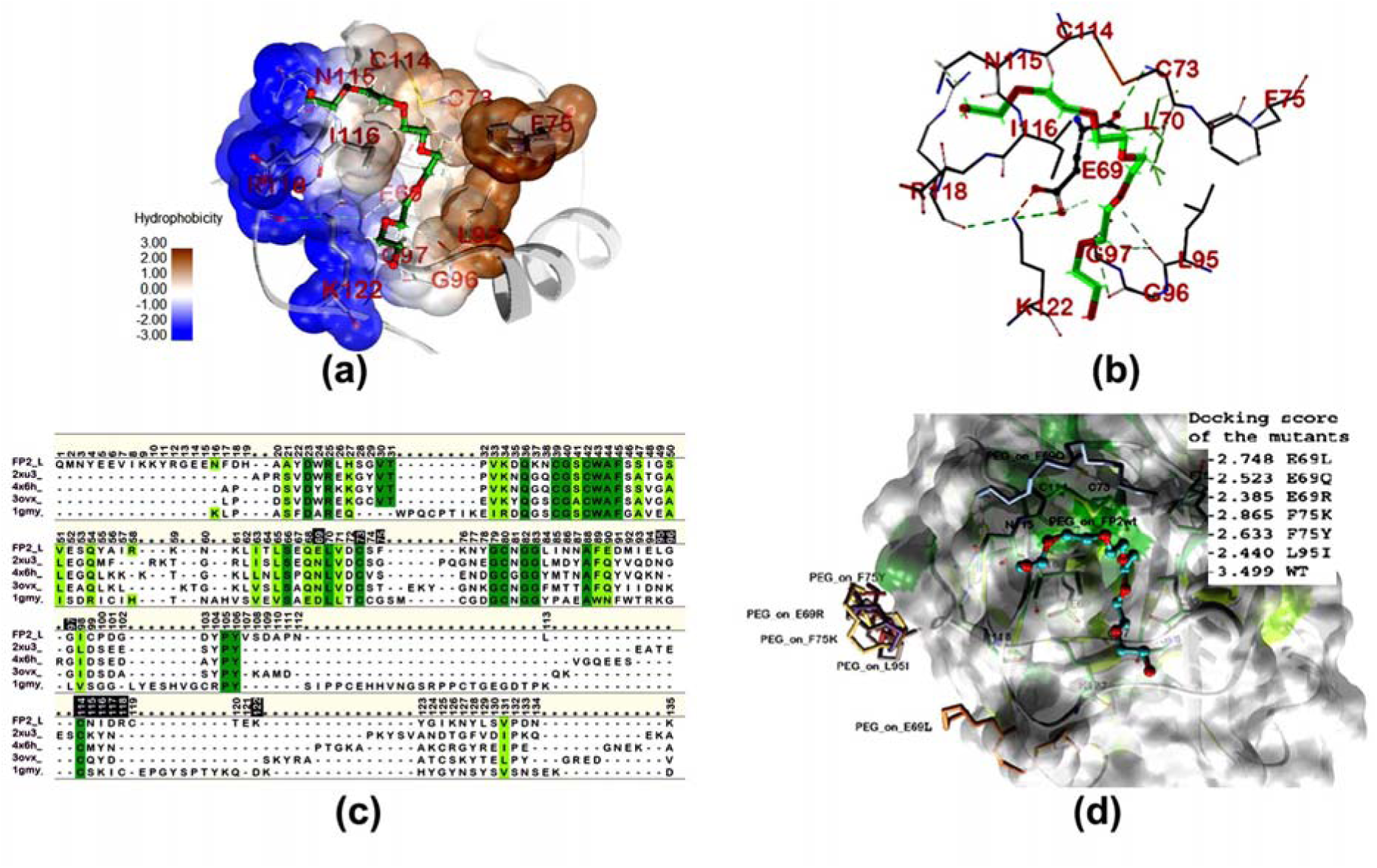
Analysis of allosteric site. (a) The hydrophobicity-based surface presentation of FP2, focussing at allosteric site, showing the central part of PEG400 docked in a well-defined cleft whereas either end is solvent exposed, (b) Interaction of PEG400 with allosteric site residues of FP2 in detail, (c) Sequence alignment of FP2 on human cathepsins L, K, S and B based on structural superposition. Deep and light green colour codes represent identity and similarity of sequence respectively. The allosteric residues in FP2 are shaded black, (d) Surface presentation of 3D structure of FP2 wild-type with surface colour according to sequence conservation mentioned in (c). The PEG400 at the allosteric site is represented as ball and stick model. The relative position of top score PEG400 molecules when docked in the FP2 mutants (allosteric site residues) are shown in stick model with annotation for their cognate mutant FP2. Inset shows the docking scores for the top-ranked poses of PEG400 for wild-type and mutants.

**Figure 11.**
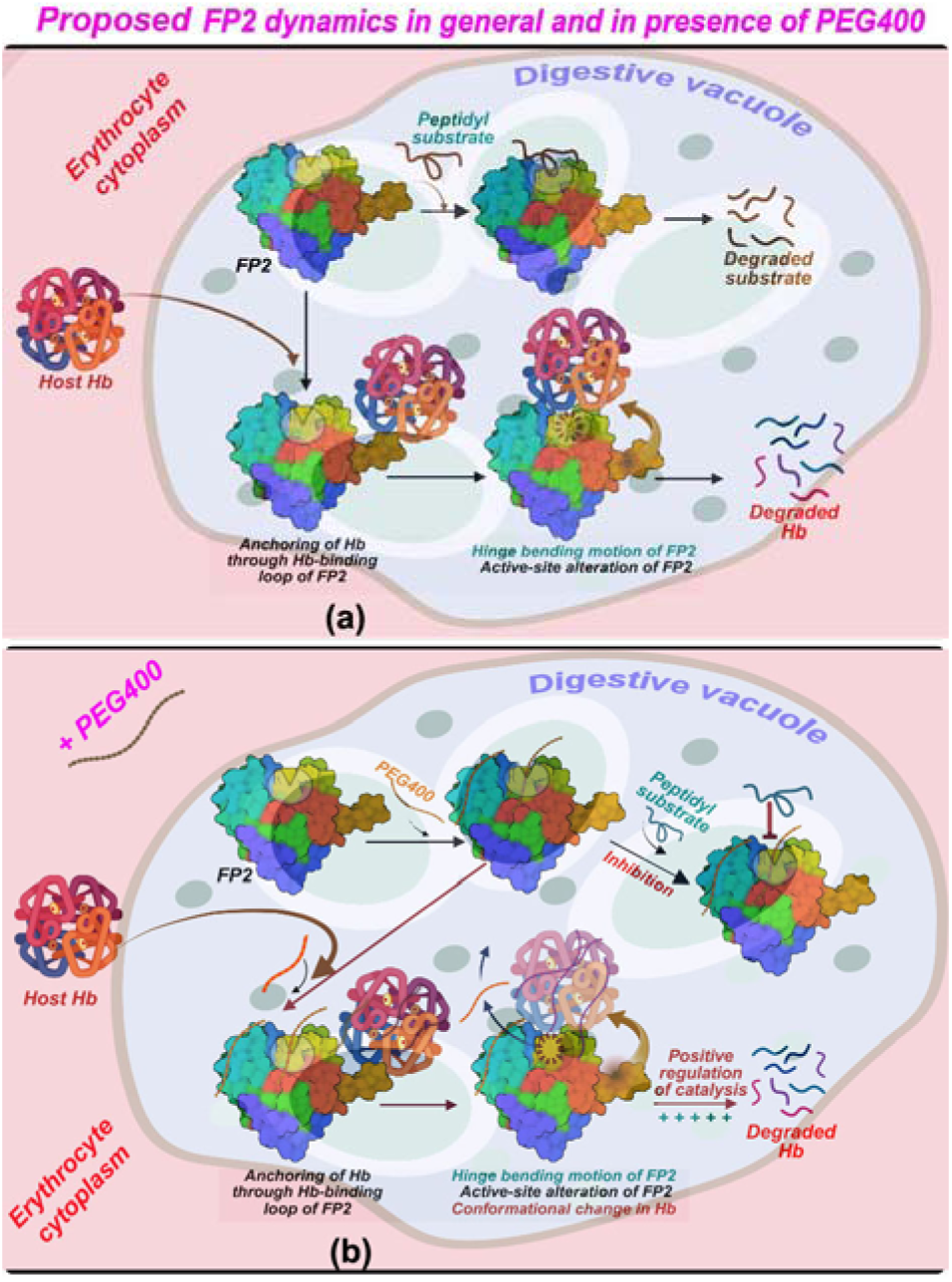
Schematic representation of FP2 activity and regulation during the trophozoite stage of plasmodium, a) in general, b) The PEG 400 induced regulation of activity of FP2 in a well-defined hierarchy.

Among these, Glu69 is important for not only providing a hydrogen-bond with PEG400 (Fig 10b), but also making h-bond/salt-bridge network with surrounding residues forming the allosteric cleft. These interactions are observed almost 21% in the dynamics trajectory (Table S5). Two hydrophobic residues Phe75 and Leu95 form the other wall of the allosteric cleft (Fig 10a, b).

To access their functional contribution, we generated 20 amino residue titration of GLu69, Phe75 and Leu95 and picked up the stabilizing and neutral mutants (Table S6) for docking with box size (20 20 20 Å) and center (-23.0 0.0 -29.0 Å), covering the full L-domain excluding the catalytic cleft. The results show none of the allosteric mutants reproduce the original PEG binding pose (Fig 10d). Moreover, all mutant dockings have reduced score compared to wild-type FP2-PEG400 (Fig 10d) complex. Even a subtle modification, such Phe75→Tyr75 substitution, alters the cavity size and nature by re-orienting nearby Leu95 (Fig 10b). This study further indicates the structural uniqueness of this site and supports its role as a potential allosteric regulator of FP2.

## DISCUSSION

### Role of Falcipain 2

Falcipains, categorized under clan C1A of papain-like cysteine proteases, are found ubiquitously across a diverse range of eukaryotes including invertebrates, plants, and human parasites [34]. The endogenous involvement and regulation of these proteases in cellular pathways still constitute an ongoing area of study. Falcipain 2 is amongst the key proteases involved in the hemoglobin degradation pathway during the intra-erythrocytic stage of *Plasmodium falciparum* infection that allows the parasite to obtain amino acids facilitating its survival and proliferation in its acidic digestive vacuole. Apart from its role as a hemoglobinase, it also cleaves skeletal membrane proteins of iRBCs at late trophozoite/schizont stages causing membranein stability to ease the iRBC rupture and merozoite egress [35]. Hemoglobin degradation and detoxification of free heme by crystalline hemozoin formation occur sequentially and are regulated spatio-temporally. Till date, majority of anti-malarial drugs have tried to exploit this seminally vulnerable pathway in *Plasmodium* life cycle. Therefore, comprehending the entirety of this process and that of the individual enzymes, is important to combat the disease, especially in the light of emerging drug resistance. Our basic objective of this study was two-fold; primarily to understand the regulatory dynamics of FP2 activity from a structural perspective in general and subsequently to access if the regulation could be potentially modulated in presence of any ligand.

### PEG400 as regulator of FP2 activity

Till date, no reports have indicated that the water-soluble polymer PEG400 inhibits the general proteolytic activity of *Plasmodial*FP2. Our study has shown that PEG has a multifaceted regulatory role in promoting FP2 activity. Thus, the outcome of the study provides a mechanistic basis for the development of PEG derivatives with anti-malarial potential. In this work, we have investigated in-depth mechanisms of how the hemoglobinase activity of FP2 is allosterically controlled by the binding of PEG. Despite significant advances, studying allosteric regulation poses a bottleneck due to the inherent complexity of allostery. Identifying a potential allosteric site of FP2 could fundamentally enhance our understanding of the molecular mechanisms governing this enzyme’s function.

The increased hemoglobinase activity in the presence of PEG400 implies that PEG400 may allosterically facilitate a structural re-organization in the active site, conducive to Hb proteolysis. Since structural change and delocalization of heme moiety in Hb have also been observed in presence of PEG400, one cannot exclude the possibility of either individual or combined structural change(s) brought about in the hemoglobin molecule making it more susceptible for proteolysis. In this context, it is worth mentioning that the collagenase activity of human cathepsin K during ECM remodeling is allosterically controlled by glucosaminoglycan [14]. An extrapolation of our study envisages a similar regulation of FP2 for its hemoglobinase activity *in-vivo*.

Here we have shown that in presence of PEG400, FP2 activity is unidirectional for degrading intact Hb through suppressing small peptide degradation. Our findings unveil a unique molecular mechanism governing the regulation of FP2 in *Plasmodium falciparum* food vacuole through mimicking thepresence of PEG400, *in-vitro*, in the intracellular milieu. Our findings emphasize upon the uniqueness of several activating as well as inhibitory allosteric mechanisms exhibited on FP2 that already shares many catalytic and regulatory features with human cysteine cathepsins. PEG400 inhibition mechanism of FP2, reported in this study, is distinctive owing to PEG400’s binding nature not being categorized under any of the allosteric, orthosteric or dualsteric/bitopic inhibition types. The proposed mechanism shows two different molecules of PEG400 binding separately at the orthosteric and allosteric sites of FP2, each with different binding affinities. This insight is valuable to design a dualsteric inhibitor with stringent specificity for FP2.

### Mechanistic insight of PEG400 regulation

Tracing back the basis of our study on the orientation of PEG molecules along the orthosteric and allosteric sites in the crystal structure of FP2 [6], biochemically recognizing and validating a potential allosteric modifier activity of PEG400 on FP2 functionality plausibly adds a new direction in malarial chemotherapeutic pursuits. Contextually, the molecular modeling study highlights the tentative structural positioning of the FP2 active site, its Hb-binding site and PEG-interacting sites, all in the proximity of each other, to facilitate the hemoglobinase activity of FP2 while consequently curtailing its general proteolytic activity. These findings pave the way for a novel therapeutic strategy in malaria treatment. Traditionally, PEG is employed to improve the pharmacokinetics of drug molecules, enhancing their hydrophilicity and passage of pharmacophores towards therapeutic targets. This study, however, proposes a bifunctional approach towards including PEG not only as molecule/s to boost the hydrophilicity, pharmacokinetics and passage of pharmacophores towards therapeutic targets but also as an alternant by interacting directly with a protease-based drug target to bring about its structural reorganization. The identification of an allosteric site on FP2 that is responsive to PEG binding provides a foundation for designing PEG-based or PEG-like molecules that can act as allosteric modulators.

Based on our structural model and experimental data, we have proposed a conceptual framework showing the mechanistic insight into FP2 regulation in presence of PEG [Fig. (11)]. The schematic representation provides valuable insights into the allosteric regulation of FP2. The results support the potential of PEG mimetics, as therapeutics and provide a structural and dynamic framework for exploring the antimalarial chemotherapeutic space, simultaneously acknowledging certain limitations and areas for further investigation.

Future validation will require site-directed mutagenesis of the predicted allosteric residues and proteomic profiling of haemoglobin degradation intermediates. Advanced structural approaches such as microsecond-scale simulation, cryo-EM or NMR spectroscopy could directly visualize FP2-hemogloglobin dynamics and provide stronger evidence for the proposed mechanism

### Conclusion

The identification of the allosteric site in this study provides new perspectives on understanding the functional regulation of FP2. In wider viewpoint, this site might represent a general regulatory mechanism that evolved to control the C1 family of protease which are highly homologous in nature. This is crucial for FP2, as the human genome contains 11 cysteine cathepsins with highly conserved active site structures, making off-target effects of FP2 directed inhibitors and poses the major bottleneck for considering FP2 as potential drug target for antimalarials. Interestingly our study shows the residues of the allosteric site are less conserved amongst human cathepsins and therefore demonstrates the uniqueness of the site, allowing specificity of allosteric modulator(s) towards FP2, an important host hemoglobinase of the life-cycle of the parasite. This study further endorses the hypothesis that for a flexible protein that presents multiple conformational states, either by conformational selection or upon ligand binding, may dominate the functional diversity depending on the ligand affinity and the protein conformational transition rate. As catalytic sites of cysteine proteases are highly conserved, allosteric modulation of FP2 offers a promising and safer therapeutic avenue. These findings not only expand current understanding of FP2 regulation but also establish a conceptual framework for developing next-generation antimalarials based on allosteric inhibition. In summary, the study found a unique allosteric drug target. The uniqueness comes from its non-conserved residues, making it distinct from similar host enzymes (cysteine cathepsins). The amphipathic nature of the cleft [Fig 10a] is particularly interesting for drug design as a drug molecule could be designed to interact favourably with both the polar and non-polar parts of the cleft, leading to a stronger and more specific binding.

## MATERIAL AND METHODS

### Expression and Purification of FP2 and STFA

FP2 from the *Plasmodium falciparum* 3D7 strain, cloned into the pRSETA vector, was expressed as in *E. coli* BL21 (DE3) as inclusion bodies. These were subsequently solubilized, purified, and refolded into their active form following a well-established protocol developed in our laboratory [6]. Human stefin A (STFA), cloned into the pET28a+ vector, was also expressed into *E. coli* BL21 (DE3) and purified following a standardized method previously established in our group [12].

### Inhibition Kinetics

The potential inhibitory effects of PEG and other compounds on the enzymatic activity of FP2 were assessed using the chromogenic substrate D-VLK-pNA (Sigma-Aldrich, USA). Initial reaction velocities (VD) were determined by continuously monitoring the release of p-nitroaniline (pNA) at 410 nm, using an extinction coefficient of 8800 MD¹ cmD¹ for liberated pNA [36] with a Thermo Evolution 201 UV– visible spectrophotometer (Schaumburg, USA). Enzyme activation was optimized according to protocols previously established in our laboratory [37]. The concentration of active enzyme was determined via active-site titration using the irreversible inhibitor E-64, which binds in a 1:1 molar ratio with the active sites of cysteine proteases [38].

Kinetic parameters (K_m_ and V_max_) were derived from initial protease-substrate reactions and calculated through nonlinear regression fitting of progress curves to the Michaelis–Menten model using GraphPad Prism 6.0 (http://www.graphpad.com/prism). ICDD values, defined as the inhibitor concentration that reduces enzyme activity by 50%, were determined by plotting percent proteolytic activity against inhibitor concentration.

To calculate the inhibition constant (K_i_), the protease was incubated with two sets of inhibitor concentrations, and reaction velocities were recorded across a range of substrate concentrations. The mode of enzyme inhibition was analyzed using Lineweaver–Burk and Dixon plots. In the Lineweaver– Burk plot, 1/VD was plotted against 1/[S], while in the Dixon plot, 1/VD was plotted against 1/[I], using Origin software (OriginLab, Northampton, MA, USA). All experiments were performed in triplicates, and data were fitted to nonlinear regression models representing competitive and mixed inhibition using GraphPad Prism 6.0.

### Hemoglobinase activity of FP2

FP2 was incubated with hemoglobin (HiMedia, India) at 37 °C for 15–25 minutes, as previously described [37]. The reactions were terminated by the addition of the cysteine protease inhibitor E-64. Samples were then mixed with denaturing SDS-PAGE loading buffer, heated at 95 °C for 10 minutes, and subjected to SDS-PAGE analysis. Hemoglobin degradation was assessed by evaluating the reduction in band intensities. Gel images were processed and quantified using NIH ImageJ software (http://imagej.nih.gov/ij/). We performed SDS-PAGE-based time-course profiling of Hb degradation. Experimental parameters, including incubation time, temperature and molar ratio of FP2 to Hb were optimized to achieve complete Hb degradation.

### Azocasein assay

The proteolytic activity of FP2 in the presence of PEG was evaluated using the azocasein assay. Hydrolysis of azocasein by FP2 was monitored by measuring the increase in absorbance at 366 nm over time in 100 mM sodium acetate buffer (pH 6.5). The assay was initiated by adding 10 µl of activated FP2 to 300 µl of reaction mixture containing 0.1% azocasein and 2 mM DTT. The mixture was incubated at 37 °C for 2 hours, after which the reaction was terminated by the addition of 15 µl of ice-cold 100% trichloroacetic acid. Azo-peptide release was quantified by measuring absorbance at 366 nm using a Thermo Evolution 201 UV–visible spectrophotometer (Schaumburg, USA). A specific absorption coefficient of A^1%^_366_ = 40 was used for quantification of the azocasein degradation products [39].

### Spectroscopic analysis

Conformational changes and ligand interactions involving FP2, hemoglobin (Hb), and the FP2–Hb complex were analyzed through tryptophan fluorescence spectroscopy using a Horiba QM-8075-11-C spectrofluorometer (Kyoto, Japan). All measurements were performed at room temperature. Emission spectra were recorded from 305 to 400 nm with excitation at 195 nm. Both excitation and emission slit widths were set to 5 nm. Spectra were obtained for FP2, Hb, the FP2–Hb complex, and after incubation with PEG for 10 minutes at room temperature. We measured the intrinsic tryptophan fluorescence quenching of FP2 by recording the fluorescence intensity (FI) at 349.5 mm (λ_max_) over increasing concentration of PEG 400 to determine its K_D_. The normalized FI were plotted and the K_D_ value was obtained by nonlinear regression fitting of the resulting saturation curve.

Secondary structure analysis was conducted using far-UV circular dichroism (CD) spectroscopy on a JASCO J-815 spectropolarimeter (Easton, USA). Measurements were carried out at 20 °C in 50 mM Tris buffer (pH 8.0) using a 0.1 cm path length cuvette and a scan rate of 100 nm/min. The bandwidth was set to 2 nm, and each spectrum represented the average of three scans recorded from 200 to 260 nm at 1 nm intervals. Background spectra (buffer only) were subtracted, and ellipticity values were reported in millidegrees (mdeg). Data were collected for individual proteins, the FP2–Hb complex, and PEG-incubated samples (10 minutes at 20 °C).

To assess the environment surrounding the heme group, absorption spectra in the Soret region were recorded using a Thermo Evolution 201 UV–visible spectrophotometer (Schaumburg, USA). Samples were prepared in 50 mM Tris buffer (pH 8.0) at room temperature. Spectra were acquired from 350 to 550 nm at 1 nm intervals for Hb, the FP2–Hb complex, and after incubation with PEG for 10 minutes.

### Docking, Molecular mechanics, NMA and analysis

Docking studies were performed using individual crystal structures of FP2 (PDB ID: 7EI0), hemoglobin (PDB ID: 2DN1), and a previously reported model of the FP2–Hb complex [6] with hexa-ethylene glycol (MW 282 Da; taken from RCSB PDB databank with chemical ID P6G) mimicking PEG400. All structures were preprocessed and submitted to the AutoDock Vina docking engine [31] via the SwissDock web server [40]. Structural superposition and visualization were carried out using Discovery Studio 2022 (BIOVIA, Dassault Systèmes. San Diego, CA, USA). Potential allosteric binding sites in FP2 were identified using the CavityPlus web server (http://www.pkumdl.cn/cavityplus/) [41].

Three dimensional structural optimizations were performed in Discovery Studio 2022 using FP2-PEG complex structures having top scored poses of PEG from docking studies, each at orthosteric and allosteric sites. All complexes were parameterized using the CHARMm force field [42,43]. Structures were initially relaxed through 200 cycles of energy minimization using smart minimizer module of Discovery studio. The minimized complexes were solvated in an orthorhombic box containing TIP3P water molecules and counter ions, ensuring a minimum distance of 7 Å between protein atoms and the box edge The solvated complexes were further energy minimized for 200 cycles under harmonic restraints on backbone atoms allowing side chains to relax, followed by full minimization of all atoms for 1500 cycles using the conjugate gradient algorithm resulting in a RMS gradient of ∼0.07 Kcal/mol/Å. The interaction energies were calculated on the minimized complexes. Allosteric communication pathways were analyzed on the minimized structure using the Ohm server (https://dokhlab.med.psu.edu/ohm/), which predicts residue networks mediating remote effects.

Normal mode analysis (NMA) was conducted on the FP2–Hb complex and its PEG-bound structure using the ElNémo web server [44] using default parameters.

A short simulation was carried out on FP2 minimized structure using a heating phase from 50 K to 300 K for 100 ps, equilibration at 300 K for 500 ps, and a 100 ns production run using a 2 fs time step and a 20 ps saving interval (total 5000 frames were collected). Generalized Born (GB) implicit solvent model was used and The Leapfrog Verlet method was followed keeping the short-range nonbonded interaction cut-off at 12 Å, in a NPT production ensemble. Bond constraint was applied for all H atoms using the SHAKE algorithm implemented in the program. Trajectory analysis was performed to evaluate stability, including assessment of potential energy, interaction energy, root-mean-square deviation (RMSD), and root-mean-square fluctuation (RMSF) for individual residues.

PCA and cross correlation matrix were computed with ProDy [45] using default settings. To represent the movement directions captured by the eigenvectors, the porcupine plot was generated using VMD software [46].

*In silico* mutational analyses were performed using Biovia Discovery Studio (Dassault Systèmes. San Diego, CA, USA) on Glu69, Phe75 and Leu95 taking into consideration the potential impacts of protein ionization, solution pH, ionic strength, and temperature-dependent effects. Three residues were mutated to all 20 amino acid combinations and results were analysed for stability of mutant(s). The neutral and stable mutants were selected for docking analysis as described earlier.

## Abbreviation

PEG: polyethylene glycol;
FP2: falcipain 2;
Hb: hemoglobin;
ACT: Artemisinin-based combination therapies;
CD: circular dichroism;
NMA: normal mode analysis;
MD: molecular dynamics

## ACKNOWLEDGEMENT

We are grateful for the intramural research grant, provided by Saha Institute of Nuclear Physics (SINP) under the Department of Atomic Energy (DAE), Government of India. Authors acknowledge the central instrumental facility of Bose Institute, Kolkata where CD experiments were performed.

## AUTHOR CONTRIBUTIONS

Conception and design of the study was by SB. BN and SB performed the experiments and data analysis. BN, SC and SB contributed writing of the manuscript.

Supplementary documents file Supplementary.docx

## References

1. World Health Organization (WHO) (2023) World malaria report 2023. WHO, Geneva, Switzerland.

2. Tse EG, Korsik M & Todd MH (2019) The past, present and future of anti-malarial medicines. Malar J 18, 93.

3. Pandey KC, Wang SX, Sijwali PS, Lau AL, McKerrow JH & Rosenthal PJ (2005) The Plasmodium falciparum cysteine protease falcipain-2 captures its substrate, hemoglobin, via a unique motif. Proc Natl Acad Sci USA 102, 9138–9143.

4. Subramanian S, Hardt M, Choe Y, Niles RK, Johansen EB, Legac J, Gut J, Kerr ID, Craik CS & Rosenthal PJ (2009) Hemoglobin cleavage site-specificity of the Plasmodium falciparum cysteine proteases falcipain-2 and falcipain-3. PLoS One 4, e5156.

5. Ramjee MK, Flinn NS, Pemberton TP, Quibell M, Wang Y & Watts JP (2006) Substrate mapping and inhibitor profiling of falcipain-2, falcipain-3 and berghepain-2: implications for peptidase anti-malarial drug discovery. Biochem J 399, 47–57.

6. Chakraborty S, Alam B & Biswas S (2022) New insights of falcipain-2 structure from Plasmodium falciparum 3D7 strain. Biochem Biophys Res Commun 590, 145–151.

7. Subramanian S, Hardt M, Choe Y, Niles RK, Johansen EB, Legac J, et al. Hemoglobin cleavage site-specificity of the Plasmodium falciparum cysteine proteases falcipain-2 and falcipain-3. PLoS One. 2009;4:e5156.

8. Pandey KC, Wang SX, Sijwali PS, Lau AL, McKerrow JH, Peterson TC, et al. The Plasmodium falciparum cysteine protease falcipain-2 captures its substrate, hemoglobin, via a unique motif. Proc Natl Acad Sci USA. 2005;102:9138–9143.

9. Pasupureddy R, Verma S, Pant A, Sharma R, Seshadri S, Pande V, et al. Crucial residues in falcipains that mediate hemoglobin hydrolysis. Exp Parasitol. 2019;197:43–50.

10. Raphael P, Takakuwa Y, Manno S, Liu SC, Chishti AH & Hanspal M (2000) A cysteine protease activity from Plasmodium falciparum cleaves human erythrocyte ankyrin. Mol Biochem Parasitol 110, 259–272.

11. Glushakova S, Mazar J, Hohmann-Marriott MF, Hama E & Zimmerberg J (2009) Irreversible effect of cysteine protease inhibitors on the release of malaria parasites from infected erythrocytes. Cell Microbiol 11, 95–105.

12. Chakraborty S & Biswas S (2023) Structure-based optimization of protease-inhibitor interactions to enhance specificity of human Stefin-A against falcipain-2 from the Plasmodium falciparum 3D7 strain. Biochemistry 62, 1053–1069.

13. Pandey KC & Dixit R (2012) Structure-function of falcipains: malarial cysteine proteases. J Trop Med 2012, 345195.

14. Novinec M, Korenč M, Caflisch A, Ranganathan R, Lenarčič B & Baici A (2014) A novel allosteric mechanism in the cysteine peptidase cathepsin K discovered by computational methods. Nat Commun 5, 3287.

15. Feig M & Sugita Y (2012) Variable interactions between protein crowders and biomolecular solutes are important in understanding cellular crowding. J Phys Chem B 116, 599–605.

16. Gnutt D, Gao M, Brylski O, Heyden M & Ebbinghaus S (2015) Excluded-volume effects in living cells. Angew Chem Int Ed Engl 54, 2548–2551.

17. Biswas S, Kundu J, Mukherjee SK & Chowdhury PK (2018) Mixed macromolecular crowding: A protein and solvent perspective. ACS Omega 3, 4316–4330.

18. Parray ZA, Ahmad F, Alajmi MF, Hussain A, Hassan MI & Islam A (2021) Interaction of polyethylene glycol with cytochrome c investigated via in vitro and in silico approaches. Sci Rep 11, 6475.

19. Pastor I, Pitulice L, Balcells C, Vilaseca E, Madurga S, Isvoran A, Cascante M & Mas F (2014) Effect of crowding by Dextrans in enzymatic reactions. Biophys Chem 185, 8–13.

20. Homchaudhuri L, Sarma N & Swaminathan R (2006) Effect of crowding by dextrans and Ficolls on the rate of alkaline phosphatase–catalyzed hydrolysis: A size-dependent investigation. Biopolymers 83, 477–486.

21. Pant A, Kumar R, Wani NA, Verma S, Sharma R, Pande V, Saxena AK, Dixit R, Rai R & Pandey KC (2018) Allosteric site inhibitor disrupting auto-processing of malarial cysteine proteases. Sci Rep 8, 16193.

22. Hernández González JE, Salas-Sarduy E, Hernández Alvarez L, Barreto Gomes DE, Pascutti PG, Oostenbrink C & Leite VBP (2021) In silico identification of noncompetitive inhibitors targeting an uncharacterized allosteric site of falcipain-2. J Comput Aided Mol Des 35, 1067–1079.

23. Marques AF, Gomes PSFC, Oliveira PL, Rosenthal PJ, Pascutti PG & Lima LMTR (2015) Allosteric regulation of the Plasmodium falciparum cysteine protease falcipain-2 by heme. Arch Biochem Biophys 573, 92–99.

24. Novinec M, Rebernik M & Lenarčič B (2016) An allosteric site enables fine-tuning of cathepsin K by diverse effectors. FEBS Lett 590, 4507–4518.

25. Li Z, Yasuda Y, Li W, Bogyo M, Katz N, Gordon RE, Fields GB & Brömme D (2004) Regulation of collagenase activities of human cathepsins by glycosaminoglycans. J Biol Chem 279, 5470–5479.

26. Almeida PC, Nantes IL, Chagas JR, Rizzi CC, Faljoni-Alario A, Carmona E, Juliano L, Nader HB & Tersariol IL (2001) Cathepsin B activity regulation. Heparin-like glycosaminoglycans protect human cathepsin B from alkaline pH-induced inactivation. J Biol Chem 276, 944–951.

27. Hernández González JE, Alberca LN, Masforrol González Y, Reyes Acosta O, Talevi A & Salas-Sarduy E (2022) Tetracycline derivatives inhibit plasmodial cysteine protease falcipain-2 through binding to a distal allosteric site. J Chem Inf Model 62, 159–175.

28. Okeke CJ, Musyoka TM, Sheik Amamuddy O, Barozi V & Tastan Bishop Ö (2021) Allosteric pockets and dynamic residue network hubs of falcipain 2 in mutations including those linked to artemisinin resistance. Comput Struct Biotechnol J 19, 5647–5666.

29. Almeida PC, Nantes IL, Rizzi CC, Júdice WA, Chagas JR, Juliano L, Nader HB & Tersariol IL (1999) Cysteine proteinase activity regulation. A possible role of heparin and heparin-like glycosaminoglycans. J Biol Chem 274, 30433–30438.

30. Roy S, Das Chakraborty S & Biswas S (2018) Not all pycnodysostosis-related mutants of human cathepsin K are inactive – crystal structure and biochemical studies of an active mutant I249T. FEBS J 285, 4265–4280.

31. Eberhardt J, Santos-Martins D, Tillack AF & Forli S (2021) AutoDock Vina 1.2.0: New docking methods, expanded force field, and Python bindings. J Chem Inf Model 61, 3891–3898.

32. Huang W, Lu S, Huang Z, Liu X, Mou L, Luo Y, Zhao Y, Liu Y, Chen Z, Hou T & Zhang J (2013) Allosite: a method for predicting allosteric sites. Bioinformatics 29, 2357–2359.

33. Shamsi A, Mohammad T, Anwar MA, Alajmi MF, Hussain A, Islam A, et al. Insight into the binding of PEG-400 with eye protein alpha-crystallin: biophysical and computational approaches. J Biomol Struct Dyn. 2021;39:3847–3862.

34. Brocklehurst K, Willenbrock F & Salih E (1987) Cysteine proteinases. In Hydrolytic Enzymes: New Comprehensive Biochemistry (Neuberger A & Brocklehurst K, eds), Vol. 16, pp. 39–158. Elsevier, Amsterdam, The Netherlands.

35. Hanspal M, Dua M, Takakuwa Y, Chishti AH & Mizuno A (2002) Plasmodium falciparum cysteine protease falcipain-2 cleaves erythrocyte membrane skeletal proteins at late stages of parasite development. Blood 100, 1048–1054.

36. Mole JE & Horton HR (1973) Kinetics of papain-catalyzed hydrolysis of N-benzoyl-L-arginine-p-nitroanilide. Biochemistry 12, 816–822.

37. Alam B & Biswas S (2019) Inhibition of Plasmodium falciparum cysteine protease falcipain-2 by a human cross-class inhibitor serpinB3: A mechanistic insight. Biochim Biophys Acta Proteins Proteom 1867, 854–865.

38. Barrett AJ, Kembhavi AA, Brown MA, Kirschke H, Knight CG, Tamai M & Hanada K (1982) L-trans-Epoxysuccinyl-leucylamido(4-guanidino)butane (E-64) and its analogues as inhibitors of cysteine proteinases including cathepsins B, H and L. Biochem J 201, 189–198.

39. Chen CY, Luo SC, Kuo CF, Lin YS, Wu JJ, Lin MT, Liu CC, Jeng WY & Chuang WJ (2003) Maturation processing and characterization of streptopain. J Biol Chem 278, 17336–17343.

40. Grosdidier A, Zoete V & Michielin O (2011) SwissDock, a protein-small molecule docking web service based on EADock DSS. Nucleic Acids Res 39, W270–W277.

41. Xu Y, Wang S, Hu Q, Gao S, Ma X, Zhang W, Shen Y, Chen F, Lai L & Pei J (2018) CavityPlus: a web server for protein cavity detection with pharmacophore modelling, allosteric site identification and covalent ligand binding ability prediction. Nucleic Acids Res 46, W374–W379.

42. Brooks BR, Brooks CL 3rd, Mackerell AD Jr, Nilsson L, Petrella RJ, Roux B, Won Y, Archontis G, Bartels C, Boresch S, et al. (2009) CHARMM: The biomolecular simulation program. J Comput Chem 30, 1545–1614.

43. Jo S, Kim T, Iyer VG & Im W (2008) CHARMM-GUI: a web-based graphical user interface for CHARMM. J Comput Chem 29, 1859–1865.

44. Suhre K & Sanejouand YH (2004) ElNemo: a normal mode web server for protein movement analysis and the generation of templates for molecular replacement. Nucleic Acids Res 32, W610–W614.

45. Bakan A, Meireles LM, Bahar I. ProDy: protein dynamics inferred from theory and experiments. Bioinformatics. 2011;27(11):1575–1577.

46. Humphrey W, Dalke A, Schulten K. VMD: Visual molecular dynamics. J Mol Graph. 1996;14(1):33–38

