## Supplementary documents for "PEG400 regulates Falcipain 2 activity through an unprecedented allosteric mechanism"

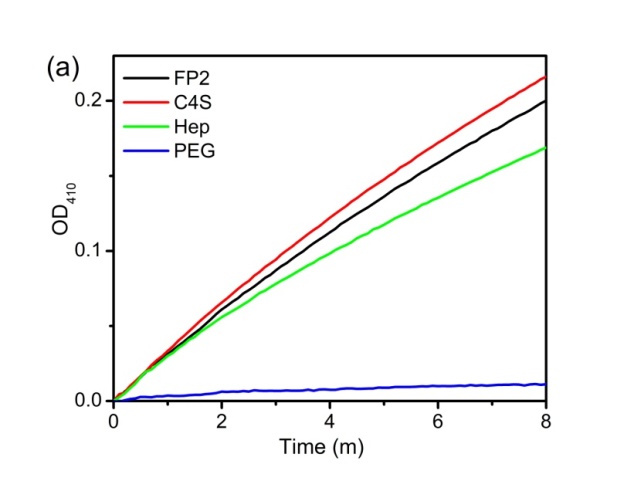

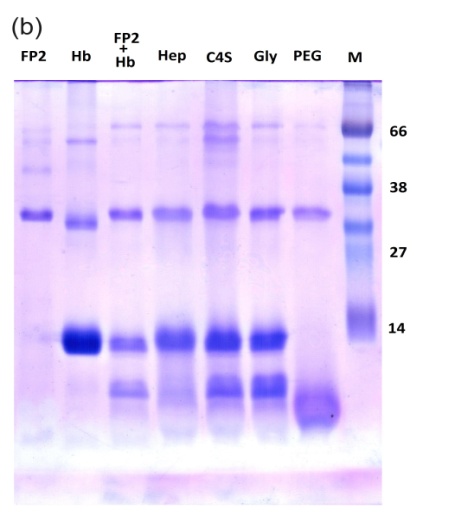

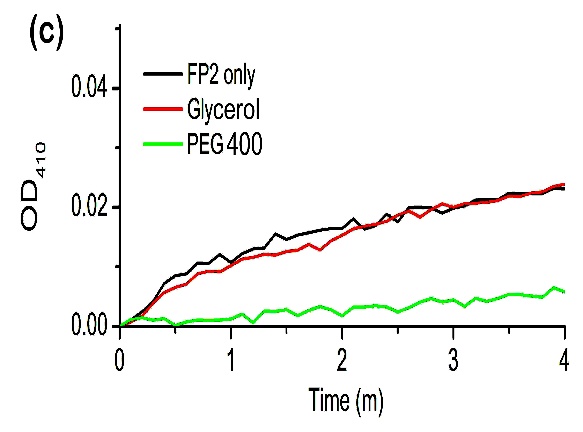

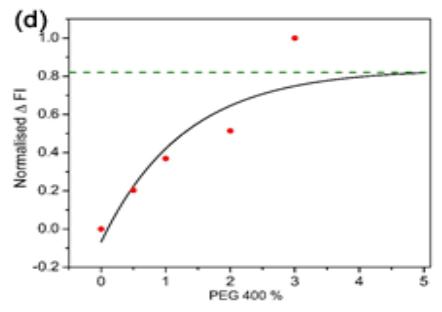

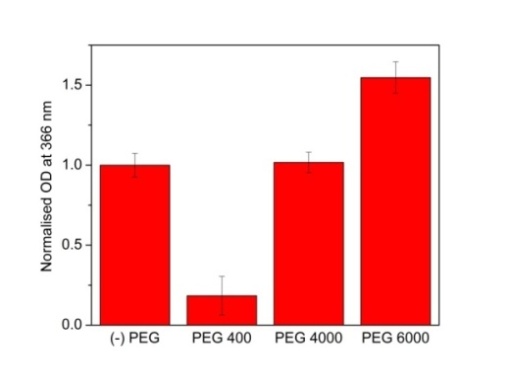

**(e)**

Fig. S1. (a) FP2 activity standard curve showing optical density (OD) at 410 nm against time using chromogenic substrate D-VLK-pNA, of untreated FP2 (denoted as ‘FP2’) and FP2 in presence of 1.5 mg/ml Chondroitin sulphate sodium salt ( ‘C4S’), 1.5 mg/ml Heparin sodium salt ( ‘Hep’) and 10 % PEG 400 ( ‘PEG’). (b) Hemoglobin (Hb) degradation analysis following 10 min incubation at 37˚C using 15 % SDS PAGE where FP2, Hb and (FP2 + Hb) denotes FP2 alone, Hb alone and a mixture of FP2 and Hb respectively. Hep, C4S, Gly and PEG denote Hb incubated withFP2 in presence of Heparin sodium salt (1.5 mg/ml), Chondroitin sulphate sodium salt (1.5 mg/ml), Glycerol (10 %) and PEG 400 (10 %) respectively. (c) FP2 activity curve showing optical density (OD) at 410 nm over time using the chromogenic substrate D-VLK-pNA for untreated FP2 (FP2 only), FP2 with 2 % glycerol (Glycerol) and 2 % PEG 400 (PEG 400). (d) Normalised intrinsic tryptophan fluorescence intensity of FP2 with increasing concentration of PEG400. KD was derived from non-linear regression fitting of the saturation curve. (e) Proteolytic activity of FP2 using azocasein assay in absence PEG [denoted as (-) PEG] and in presence of 2% PEG 400, 4000 and 6000. Hydrolysis of azocasein was determined at 366 nm by increase in OD after incubation at 37˚C for 1.5 hr in 50 mM Na-acetate pH 6.5. OD values were normalised to the activity of to FP2 without PEG.

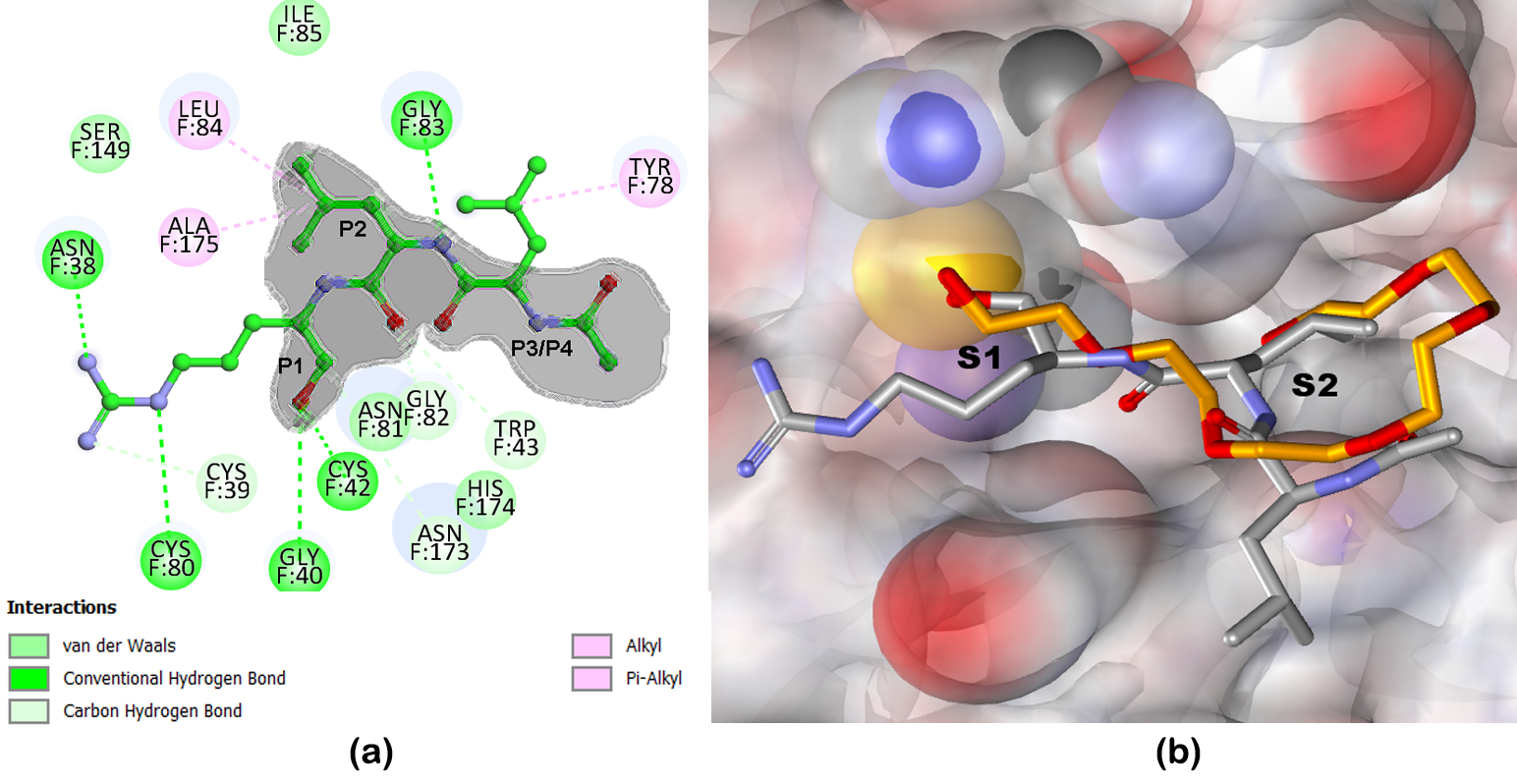

Fig. S2 (a) Interaction of Leupeptin inhibitor with catalytic cleft of FP2 in 2D interaction diagram (b) 3D diagram showing PEG400 superposed at the FP2 catalytic cleft. Grey shading in (a) highlights the common interacting regions of Leupeptin and PEG400.

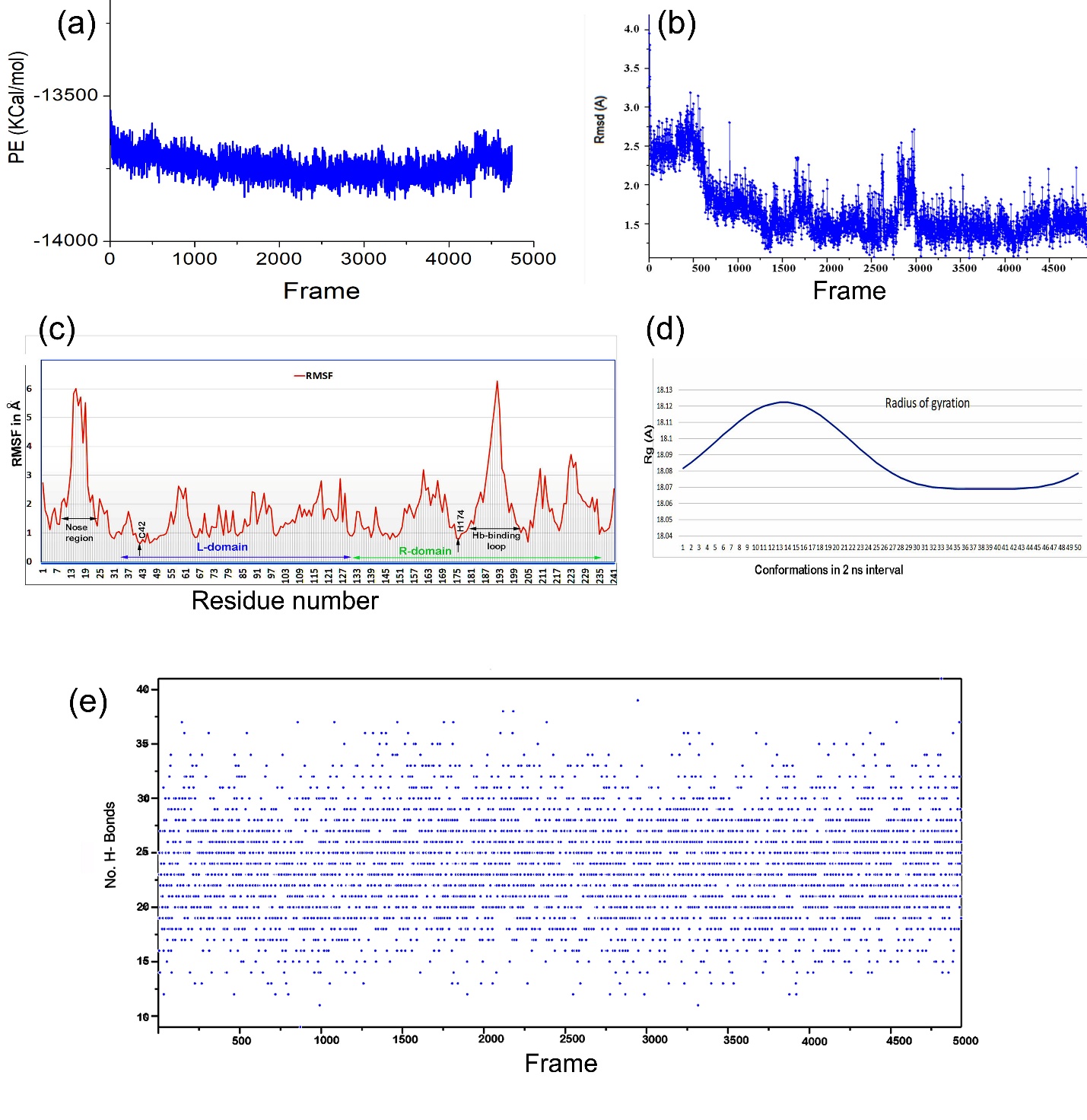

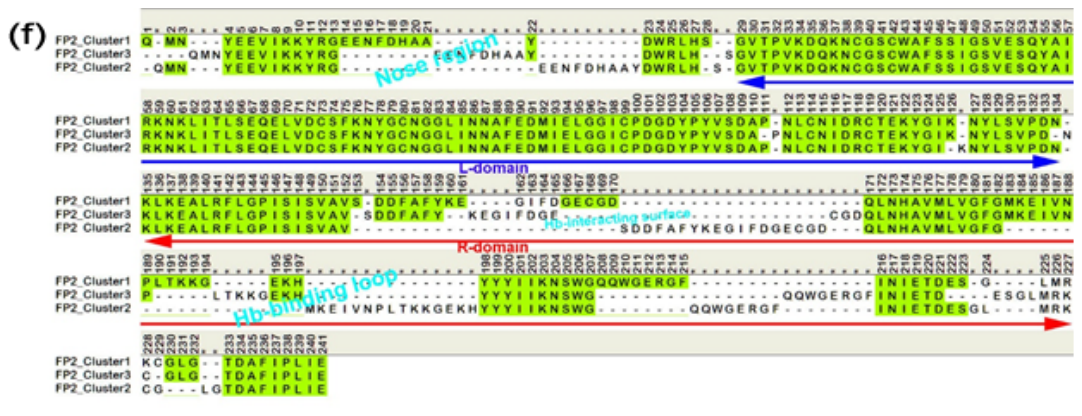

Fig S3. MD simulation analyses. a) Potential Energy (PE), b) Root mean square deviation (Rmsd), c) Root mean square fluctuation (RMSF), d) Radius of gyration (Rg) and e) Number of Hydrogen bond of the FP2 structure in the trajectory frames. f) Structural superposition of three representative structures are projected in 1D primary structure to understand region of structural deviation amongst the clusters.

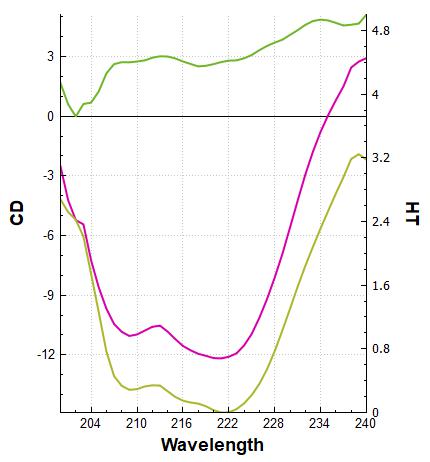

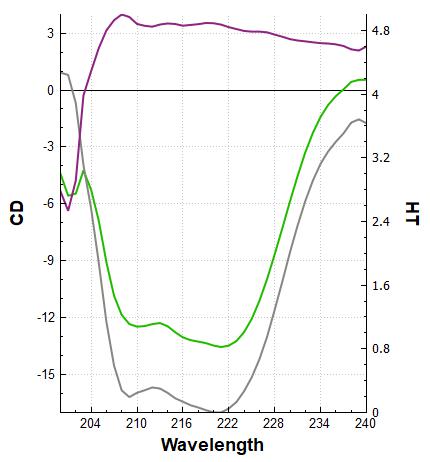

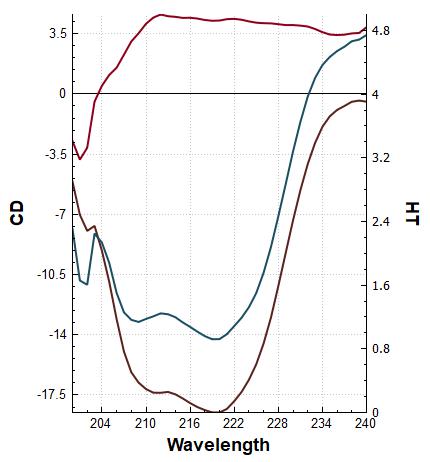

Hb	FP2-Hb	FP2

Fig. S4. Far-UV CD and difference spectra with 0 and 1% PEG400

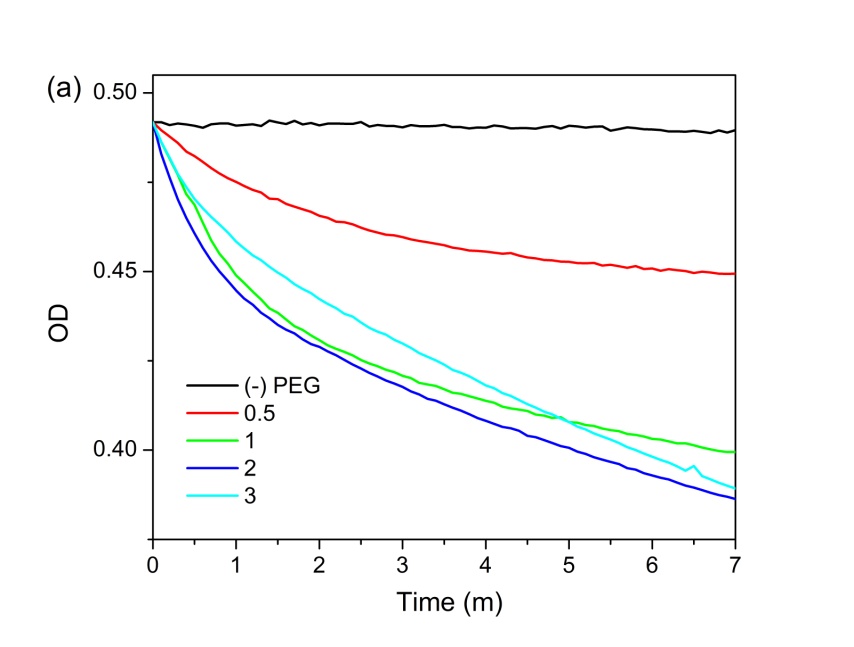

Fig. S5. OD at 406 nm as a function of time of Hb in presence of PEG400 with different concentration indicated in % (w/v).

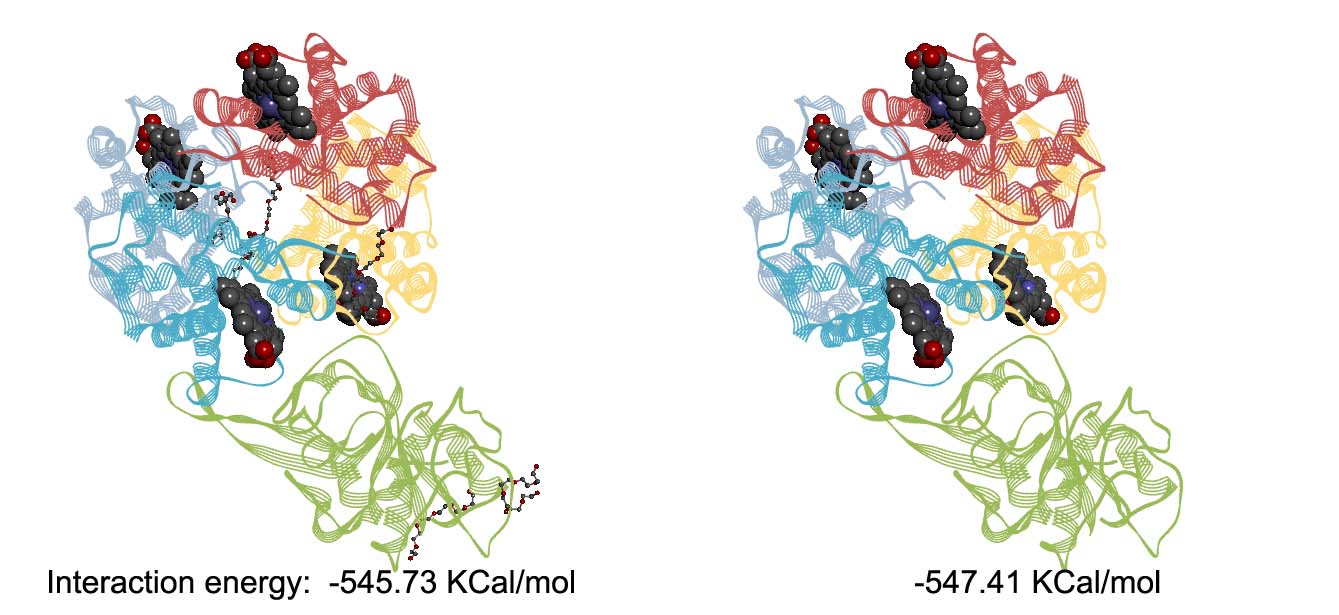

Fig. S6. FP2-Hb interaction energy in presence and absence of PEG

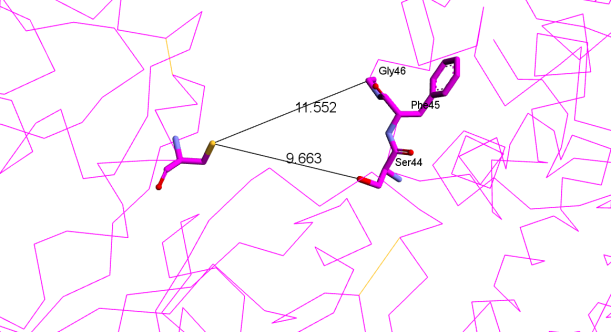

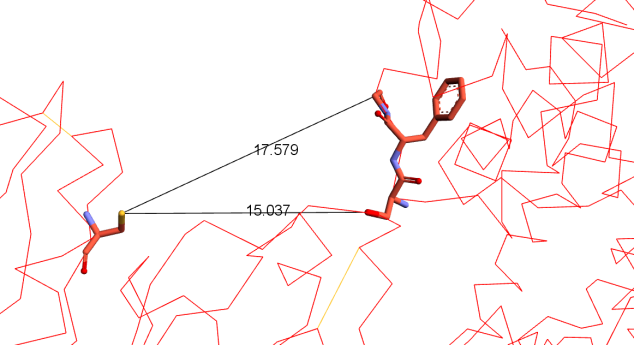

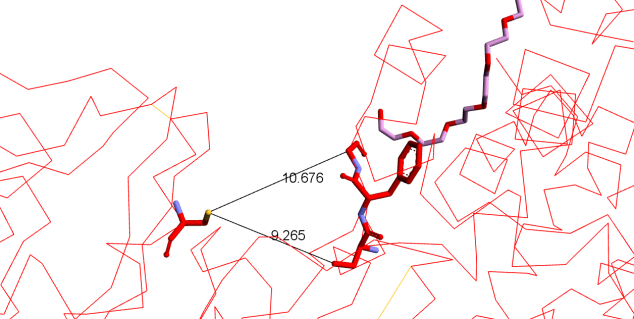

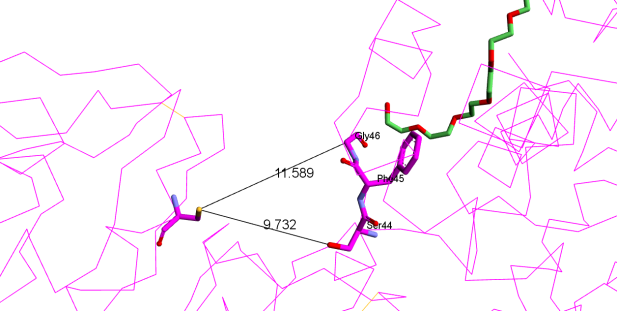

Fig. S7. FP2-Hb proximity during two extreme position in NMA in absence (upper panel) and in presence (lower panel) of PEG400.

**Table S1:** Cluster properties from PCA analysis and comparison with crystal structure of FP2

| Cluster number | Volume Å^3^ | Surface Å^2^ | Radius of gyration Å | Allosteric-site  Volume Å^3^ | Allosteric-site  Surface Å^2^ | Docking Score  (Kcal/mol) |
| --- | --- | --- | --- | --- | --- | --- |
| 1 | 34,956.2 | 11,484.8 | 18.07 | 1,347.81 | 1,198.53 | -3.498 |
| 2 | 35,055.5 | 12,313.2 | 18.08 | 1,398.45 | 1,123.01 | -2.523 |
| 3 | 35,077.4 | 12,026.5 | 18.11 | 1,382.0 | 1,129.22 | -3.025 |
| Crystal | 30,408.4 | 8,785.75 | 17.37 | 1,293.71 | 1,013.65 | -3.499 |

Table S2. Variation of fluorescence emission maxima, (λ_max_) and fluorescence intensity (FI) of tryptophan fluorescence with [PEG 400]

| [PEG]  (%) | Hb | | FP2 | | Hb + FP2 | |
| --- | --- | --- | --- | --- | --- | --- |
|  | λ_max_  (nm) | FI at 337.5 nm  (10^6^ cps) | λ_max_  (nm) | FI at 349.5 nm  (10^6^ cps) | λ_max_  (nm) | FI at 347.5 nm  (10^6^ cps) |
| 0 | 337.5 | 1.75142 | 349.5 | 4.0510 | 347.5 | 2.8414 |
| 0.5 | 338 | 1.4932 | 350.0 | 3.9185 | 345 | 3.03022 |
| 1 | 340 | 1.28569 | 350.5 | 3.8100 | 347.5 | 2.81136 |
| 2 | 339 | 1.24655 | 350.5 | 3.7155 | 347 | 2.86534 |
| 3 | 339.5 | 1.39271 | 351.5 | 3.3982 | 347 | 2.68118 |

Table S3. Variation of Absorption peak at soret region with [PEG 400]

| [PEG]  (%) | Hb | | Hb + FP2 | |
| --- | --- | --- | --- | --- |
|  | λ_max_  (nm) | OD at λ_max_ | λ_max_  (nm) | OD at λ_max_ |
| 0 | 406 | 0.52116 | 405 | 0.49837 |
| 0.5 | 411 | 0.5041 | 405 | 0.5015 |
| 1 | 411 | 0.32504 | 411 | 0.35008 |
| 2 | 415 | 0.31592 | 411 | 0.34029 |
| 3 | 416 | 0.21027 | 411 | 0.23242 |

Table S4. Pseudo-first-order rate constant of the denaturation of Hb as shown in Fig S5

| [PEG]  (%) | First phase (0-2 m) rate constant,  k_1_  (m^-1^) | Second phase (2-7 m) rate constant,  k_2_  (m^-1^) |
| --- | --- | --- |
| 0.5 | 2.656 × 10^2^ | 0.658 × 10^2^ |
| 1 | 6.563 × 10^2^ | 1.438 × 10^2^ |
| 2 | 6.275 × 10^2^ | 2.036 × 10^2^ |
| 3 | 4.942 × 10^2^ | 2.532 × 10^2^ |

Table S5: Occupancy of H-bonds involving GLU69

| **Donor** | **Acceptor** | **Occupancy** |
| --- | --- | --- |
| LYS122-Side | GLU69-Side | 1.25% |
| ARG118-Side | GLU69-Side | 1.31% |
| CYS73-Main | GLU69-Main | 5.42% |
| ASN115-Main | GLU69-Side | 0.47% |
| TYR104-Side | GLU69-Side | 0.25% |
| GLY97-Main | GLU69-Side | 8.87% |
| ASP72-Main | GLU69-Main | 0.03% |
| GLY96-Main | GLU69-Side | 1.84% |
| CYS99-Main | GLU69-Side | 0.53% |
| ILE98-Main | GLU69-Side | 0.31% |
| GLU69-Main | GLU69-Side | 0.50% |

Table S6: Twenty Single Mutations generated for FP2 Glu69, Phe75 and Leu95

| **Index** | **Mutation** | **Mutation Energy (kcal/mol)** | **Effect** |
| --- | --- | --- | --- |
| 1 | F:GLU69>GLN | -0.99 | STABILIZING |
| 2 | F:GLU69>LEU | -0.40 | NEUTRAL |
| 3 | F:GLU69>GLU | 0.10 | NEUTRAL |
| 4 | F:GLU69>ARG | 0.35 | NEUTRAL |
| 5 | F:GLU69>HIS | 1.18 | DESTABILIZING |
| 16 | F:GLU69>LYS | 3.26 | DESTABILIZING |
| 17 | F:GLU69>PRO | 3.27 | DESTABILIZING |
| 18 | F:GLU69>TYR | 4.02 | DESTABILIZING |
| 19 | F:GLU69>GLY | 4.14 | DESTABILIZING |
| 20 | F:GLU69>TRP | 18.02 | DESTABILIZING |

| **Index** | **Mutation** | **Mutation Energy (kcal/mol)** | **Effect** |
| --- | --- | --- | --- |
| 1 | F:PHE75>PHE | -0.00 | NEUTRAL |
| 2 | F:PHE75>TYR | 0.08 | NEUTRAL |
| 3 | F:PHE75>LYS | 0.39 | NEUTRAL |
| 4 | F:PHE75>LEU | 0.54 | DESTABILIZING |
| 5 | F:PHE75>ILE | 0.61 | DESTABILIZING |
| 16 | F:PHE75>SER | 1.76 | DESTABILIZING |
| 17 | F:PHE75>GLY | 2.58 | DESTABILIZING |
| 18 | F:PHE75>GLU | 3.41 | DESTABILIZING |
| 19 | F:PHE75>ASP | 3.86 | DESTABILIZING |
| 20 | F:PHE75>PRO | 4.44 | DESTABILIZING |

| **Index** | **Mutation** | **Mutation Energy (kcal/mol)** | **Effect** |
| --- | --- | --- | --- |
| 1 | F:LEU95>LEU | -0.19 | NEUTRAL |
| 2 | F:LEU95>ILE | 0.02 | NEUTRAL |
| 3 | F:LEU95>HIS | 0.62 | DESTABILIZING |
| 4 | F:LEU95>TYR | 0.64 | DESTABILIZING |
| 5 | F:LEU95>PHE | 0.69 | DESTABILIZING |
| 16 | F:LEU95>ASP | 3.39 | DESTABILIZING |
| 17 | F:LEU95>GLY | 3.96 | DESTABILIZING |
| 18 | F:LEU95>ARG | 4.01 | DESTABILIZING |
| 19 | F:LEU95>GLU | 4.63 | DESTABILIZING |
| 20 | F:LEU95>PRO | 8.52 | DESTABILIZING |
